# C5aR1 inhibition alleviates cranial radiation-induced cognitive decline

**DOI:** 10.1101/2024.07.02.601806

**Authors:** Robert P. Krattli, An H. Do, Sanad M. El-Khatib, Leila Alikhani, Mineh Markarian, Arya R. Vagadia, Manal T. Usmani, Shreya Madan, Janet E. Baulch, Richard J. Clark, Trent M. Woodruff, Andrea J. Tenner, Munjal M. Acharya

## Abstract

Cranial radiation therapy (RT) for brain cancers leads to an irreversible decline in cognitive function without an available remedy. Radiation-induced cognitive deficits (RICD) are a particularly pressing problem for the survivors of pediatric and low grade glioma (LGG) cancers who often live long post-RT lives. Radiation-induced elevated neuroinflammation and gliosis, triggered by the detrimental CNS complement cascade, lead to excessive synaptic and cognitive loss. Using intact and brain cancer-bearing mouse models, we now show that targeting anaphylatoxin complement C5a receptor (C5aR1) is neuroprotective against RICD. We used a genetic knockout, C5aR1 KO mouse, and a pharmacologic approach, employing the orally active, brain penetrant C5aR1 antagonist PMX205 to reverse RICD. Irradiated C5aR1 KO and WT mice receiving PMX205 showed significant neurocognitive improvements in object recognition memory and memory consolidation tasks. Inhibiting C5a/C5aR1 axis reduced microglial activation, astrogliosis, and synaptic loss in the irradiated brain. Importantly, C5aR1 blockage in two syngeneic, orthotopic glioblastoma-bearing mice protected against RICD without interfering with the therapeutic efficacy of RT to reduce tumor volume *in vivo*. PMX205 clinical trials with healthy individuals and amyotrophic lateral sclerosis (ALS) patients showed no toxicity, drug-related adverse events, or infections. Thus, C5aR1 inhibition is a translationally feasible approach to address RICD, an unmet medical need.

**SIGNIFICANCE:** Cranial radiotherapy for brain cancers activates CNS complement cascade, leading to cognitive decline. Ablation of the complement C5a/C5aR1 axis alleviates radiation-induced neuroinflammation, synaptic loss, and cognitive dysfunction, providing a novel tractable approach.

## INTRODUCTION

Despite being effective for tumor control, cranial radiation therapy (RT) for treating primary and metastatic brain cancers causes an unintended and irreversible impact on brain function, including cognitive impairments (1). Long-term sequelae of radiation-induced cognitive impairments (RICD) are particularly crucial for childhood brain cancer and low grade glioma (LGG) survivors who often live long post-therapy lives (2). RT triggers a CNS cascade of immune and non-immune consequences, including detrimental activation of the complement cascade (3,4), leading to long-term microglial activation and loss of synaptic integrity that eventually culminate in RICD (3). The CNS complement cascade plays unique roles in neural stem cell polarity determination (5), and pruning and maintaining synaptic landscape during brain development (6). However, under pathological conditions, including Alzheimer’s disease (AD), traumatic brain injury (TBI), and RICD, excessive complement cascade activation promotes neurodegeneration via elevated neuroinflammation (3,6). We have shown that microglia-selective knockdown of an upstream complement C1q was neuroprotective against RICD, microglial activation, astrocytic hypertrophy, synaptic loss, and anaphylatoxin C5a receptor 1 (C5aR1) expression (3). However, C1q also plays an essential role in pruning synapses in the CNS (6); thus, the ablation of the upstream complement activation node might not be the optimal therapeutic approach to prevent RICD.

Genetic knockout or pharmacologic inhibition of C5aR1 significantly reduced AD neuropathologies, including plaques, gliosis, and hippocampal-dependent cognitive dysfunction (7-9). Conversely, C5a overexpression in the brain worsened hippocampal memory (10). C5a-C5aR1 signaling inhibition using a highly selective, BBB-permeable, cyclic hexapeptide antagonist, PMX205 (11), significantly reduced microglial pro-inflammatory polarization, inflammatory gene expression, and excessive synaptic loss in the AD brain (8,9,12). Notably, an FDA-approved C5aR1 antagonist, Avacopan (Tavneos^®^), is safe, well-tolerated, and effective against autoimmune disease (13), suggesting that C5aR1 inhibition can be a viable and safe strategy against RICD. However, Tavneos showed poor brain penetration (14). PMX205 has recently completed two Phase 1a dose-escalating and repeat-daily dosing studies in healthy individuals and a Phase 1b pharmacokinetics and pharmacodynamics trials in ALS patients. These trials showed no toxicity, drug-related adverse events, or infectious disease (Australian New Zealand Clinical Trials Registry, reference ACTRN12622000927729). Interestingly, C5a signaling has been suggested to increase tumorigenesis in colorectal, gastric, and ovarian tumors and glioma stem-like proliferation (15-18). Cranial RT-induced CNS complement cascade activation can lead to long-term inflammatory injury and synaptic loss in the irradiated brain (3,6).

Therefore, tractable approaches to block pro-inflammatory C5a-C5aR1 signaling without altering the therapeutic efficacy of cranial RT for brain cancers need to be investigated. The current study provides evidence that genetic KO and pharmacological C5aR1 blockade prevent cranial RT-induced gliosis and synaptic loss and ameliorate RICD. Notably, the C5aR1 blockade did not interfere with the therapeutic efficacy of RT in two glioma models, increasing the translational significance of this strategy.

## MATERIALS AND METHODS

Detailed methods are provided in the **Supplemental Information** section.

### Animal models, cranial RT, tumor induciton and cognitive function analysis

All animals used in this study were cared for per NIH guidelines and approved by the university IACUC. 15-16 weeks old male WT (C57Bl/6, Jackson, RRID: IMSR_JAX:000664) and C5aR1 KO (kind gift, R. A. Wetsel) mice were divided into the following groups: 1) WT mice receiving 0 Gy or 9 Gy cranial RT with or without PMX205 treatment (PubChem ID 90489023) and 2) C5aR1 KO mice receiving 0 Gy or 9 Gy RT. PMX205 treatment was given as one week of daily subcutaneous injections (1 mg/kg body) concurrently with drinking water (20 µg/ml) for one month. One-month post-treatment, cognitive function testing was performed, followed by euthanasia and tissue harvesting. For the tumor-bearing model, WT male mice were divided into the following groups for the astrocytoma (CT2A-Luc) and glioblastoma (GL261) tumor types: Tumor + Vehicle, Tumor + RT, Tumor + PMX205, and Tumor + RT + PMX205. Tumor was induced by stereotaxic injection of Gl261 (RRID:CVCL_Y003) or CT2A-luc (RRID:CVCL_ZJ60) cells in caudate putamen. Tumor growth (CBCT for Gl261 and BLI for CT2A) and animal health were monitored for one-to two-month post-treatment. A cohort of mice underwent cognitive function testing at three- to four weeks post-treatment. RT was administered to the non-tumor WT and C5aR1 KO mice and the tumor groups 7-9 days post-tumor induction using a SmART+ X-ray irradiator (225 kV, 20 mA, 4.91 Gy/min). One month post-RT mice were administered cognitive function tests, including open field test (OFT), elevated plus maze (EPM), object recognition memory (ORM), object location memory (OLM), and fear extinction memory (FE). Considering the health of tumor-bearing mice, ORM was conducted 3-4 weeks post-tumor induction. OFT and EPM tasks evaluate anxiety-related behavior. ORM and OLM tasks evaluate episodic memory by measuring animal’s preference to explore or recognize novel objects (ORM) or spatial locations of objects (OLM). FE determines consolidation of previously learned memory function. Detailed animal experimentation and cognitive function testing protocols are provided in **Supplemental Information**.

### Immunofluorescence staining, laser scanning confocal microscopy, volumetric quantification, cytokine and gene expression analysis

PFA-fixed brains from each treatment group underwent dual- or triple-immunofluorescence staining (3-4 sections/brain, 6-8 brains/group) including microglial activation (CD68-IBA1), astrogliosis (GFAP), C5aR1 expression, pre- and post-synaptic marker expression (synaptophysin, Syn; and PSD95). To quantify the immunoreactivity of individual and co-labeled markers, 3D blinded deconvolution and volumetric quantification were used. Please see **Supplemental Information** for detailed staining protocols, antibodies, and *in silico* modeling approaches. Data were expressed as mean percent immunoreactivity relative to the vehicle-treated control group. For the cytokine analysis, supernatants from fresh-dissected he-mi-brains (3-6 brains/group) were assayed using sandwich ELISA panel. Commercially available nCounter Mouse Neuroinflammation Panel (Nanostring, Cat. XT-CSO-MNROI1-12) was used to evaluate neuroinflammatory gene expression.

### Statistical analysis

Statistical analyses were performed to confirm overall significance (GraphPad Prism, v10.0, RRID: SCR_002798). For the analysis of C5aR KO, PMX205 treatment, and cranial irradiation, two-way ANOVA or repeated measures ANOVA and recommended multiple comparison tests were performed. For the tumor studies, P values were derived from the Mann-Whitney *U* test or long-rank test for survival studies. All results are expressed as the mean values ± SEM. All analyses considered a value of P≤0.05 to be statistically significant.

### Data availability

All data reported will be shared by the lead contact upon a reasonable request. This article did not report the original code. Any additional information required to reanalyze the data reported in this article is available from the lead contact upon a reasonable request. The NanoString Neuroinflammation gene expression panel data is available via the NCBI Gene Expression Omnibus (GEO) archive, reference number GSE282058.

## RESULTS

### Inhibition of the C5a-C5aR1 axis ameliorates RT-induced cognitive impairments

To determine if C5aR1 inhibition impacts RICD, we tested genetic and pharmacologic approaches, including a C5aR1 knockout (C5aR1-KO) mouse model and an orally active C5aR1 antagonist, PMX205, respectively. KO mice, a kind gift from R. A. Westel, were generated by targeted gene deletion (19). CNS radiobiology studies have shown that the adult female brain is protected from acute 9 to 10 Gy cranial radiation-induced cognitive dysfunction and neuronal damage (20). Thus, as a proof of principle, we have used male mice to determine the detrimental consequences of cranial RT on the complement cascade. WT C57Bl6 and C5aR1-KO male mice were exposed to 9 Gy cranial RT with the protection of eyes and cerebellum as described in the **Methods** and **Fig. 1A**. At 48h post-RT, WT mice received PMX205 treatment (20 μg/ml, or 100 μg per day) in drinking water for 1-month. In addition, mice were injected with PMX205 (1 mg/kg, subcutaneous, SQ) for the first 7-days of the antagonist treatment. Previously, using an AD mouse model(8), we found that a combination of 1 mg/kg, SQ injection for one week, and oral treatment effectively reduced about 50% of plaques, amyloid, and gliosis. Our pharmacokinetics studies have shown about 96% bioavailability of the SQ injections of PMX205 (11). Drinking water administration of PMX205 was able to sustain CNS levels of the drug. Thus, PMX205 was administered via two routes for 7 days to target the initial acute neuroinflammatory response post-RT. To confirm the genetic KO of C5aR1, qPCR and flow cytometry analyses (**Suppl. Fig. S1**) of C5aR1 (CD88) expression in the C5aR1-KO mice brains and blood showed complete knockout (>96-98%) in the 0 and 9 Gy irradiated animals.

**FIGURE 1.**
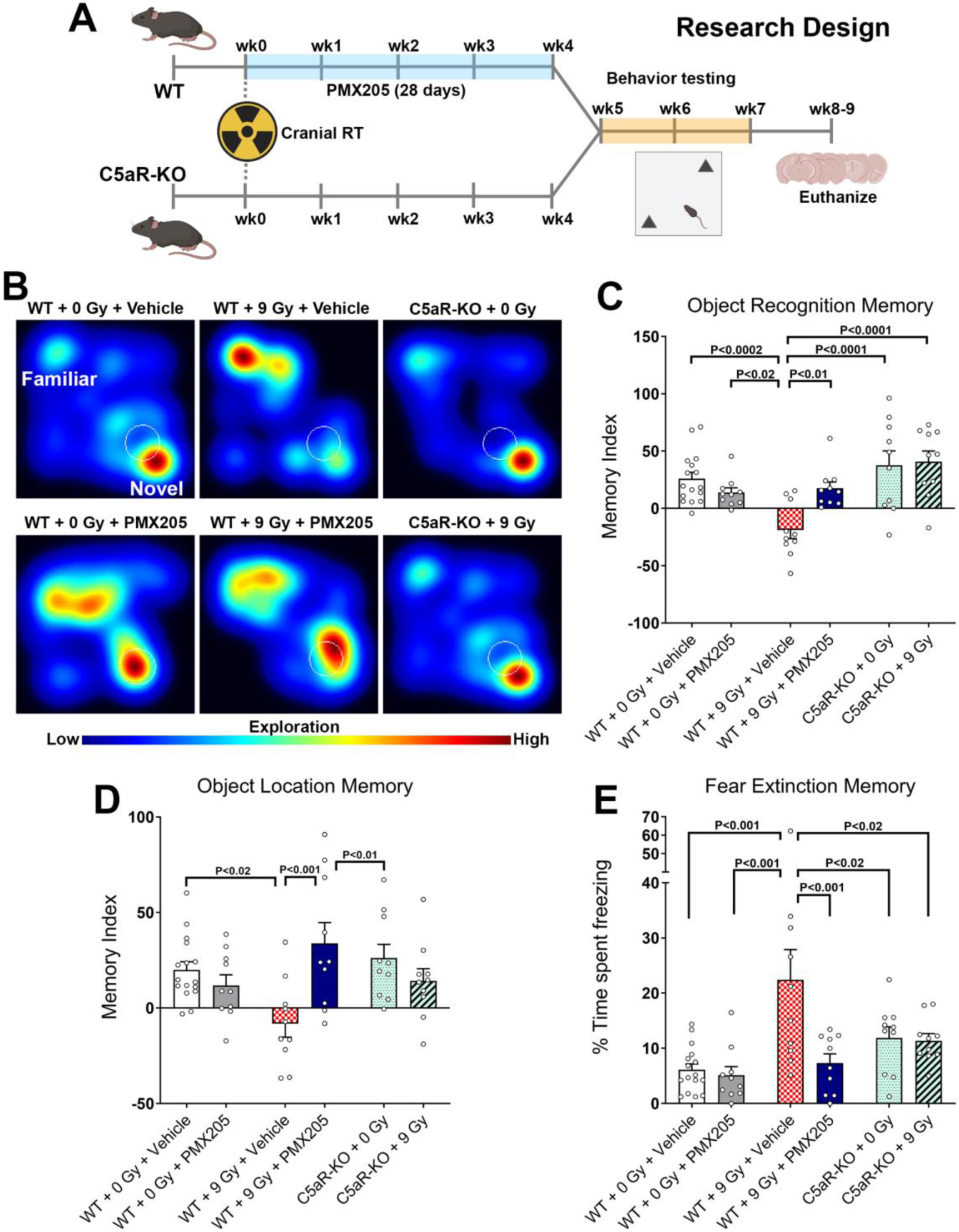
C5aR1 knockout or inhibition prevents cranial radiation-induced cognitive impairments. (A) Research design: Four-month-old C57Bl6 WT and C5aR1-KO mice received 0 or 9 Gy cranial radiotherapy (CRT). At 48h post-CRT, WT mice received PMX205 treatment (1 mg/kg SQ for the first week in addition to 20 μg/ml, or 100 μg per day in drinking water for 1 month). All mice were administered cognitive function tasks (Object Recognition Memory, ORM; Object Location Memory, OLM; and fear extinction memory consolidation, FE) 1-month later and euthanized for tissue collection. (B) Representative heat maps depicting mice exploring novel or familiar objects during the ORM task. Irradiated WT mice (WT + 9 Gy + Vehicle) did not show a preference for novel objects. (C) The tendency to explore novel objects (ORM) was derived from the Memory Index: ([Novel object exploration time/Total exploration time] – [Familiar object exploration time/Total exploration time]) ×100. WT + 9 Gy + Vehicle group showed significantly impaired cognitive function, as indicated by the reduced preference towards the novel object compared to the WT + 0 Gy + Vehicle, WT + 0 Gy + PMX205, C5aR1-KO + 0 Gy, and C5aR1-KO + 9 Gy groups. Importantly, irradiated C5aR1-KO mice (C5aR1-KO + 9 Gy) and WT + 9 Gy + PMX205 did not show a decline in the memory index. (D) Irradiated WT mice spent significantly less time exploring the novel object locations during the OLM task. WT + 9 Gy mice + PMX205 showed significantly improved memory index compared to WT + 9 Gy + Vehicle group. Irradiated C5aR1-KO mice (C5aR1-KO + 9 Gy) showed a trend of spatial exploration improvements (a positive Memory Index) during the OLM task. (E) C5aR1 inhibition or KO significantly improved fear memory consolidation (FE test). WT + 0 Gy + Vehicle and WT + 0 Gy + PMX205 showed abolished fear memory (reduced freezing, intact memory consolidation) compared with WT + 9 Gy + Vehicle mice. Importantly, WT + 9 Gy + PMX205 and C5aR1-KO + 0 Gy and 9 Gy groups successfully abolished fear memory (reduced freezing) compared to the WT + 9 Gy + Vehicle group. Mean ± SEM (*N* = 10-16 mice per group).

At one-month post-irradiation, all cohorts of mice (WT mice, 0 and 9 Gy ± PMX205; C5aR1-KO mice ± 9 Gy) were habituated and tested on the cognitive function tasks. During the open field exploration on the first day of habituation, no significant differences were found between the animal groups for the percentage of time spent in the central (open) area (30% of the total area, **Suppl. Fig. S2A**), indicating spontaneous exploration and the absence of neophobic behavior during cognitive testing. Similarly, the percentage of time spent in the open arms did not vary significantly between the groups in the elevated plus maze task (**Suppl. Fig. S2B**). After three days of habituation, animals were administered the object recognition memory (ORM) task. During the test phase, the total time exploring both familiar and novel objects was comparable between each experimental group. This behavior is apparent in the heat map of animal exploration depicting the activity for the novel or familiar objects (**Fig. 1B**). We did not find significant differences in the memory index for WT + 0 Gy + Vehicle, WT + 0 Gy + PMX205, and C5aR1-KO + 0 Gy groups indicating intact object recognition memory function. On the other hand, WT + 9 Gy + Vehicle mice showed a significant decline in the memory index (novel object exploration preference) compared to unirradiated (0 Gy) controls (WT+ 0 Gy + Vehicle, P<0.0002, **Fig. 1C**). This cognitive decline was prevented in WT + 9 Gy + PMX205 and C5aR1-KO + 9 Gy mice (**Fig. 1C**, P<0.01, P<0.0001, respectively). The performance for the object location memory (OLM) task showed an RT-induced decline in the memory index in WT + 9 Gy + Vehicle mice compared to WT + 0 Gy + Vehicle mice (P<0.02, **Fig. 1D**). PMX205 treatment of the irradiated WT mice (WT + 9 Gy + PMX205) significantly improved the memory index compared to the WT + 9 Gy + Vehicle group (P<0.01). C5aR1-KO mice receiving either 0 or 9 Gy cranial RT showed a trend for improvements in memory index, albeit it did not reach a statistical significance. Nonetheless, the behavior of 0 Gy and 9 Gy exposed C5aR1-KO mice was comparable to the 0 Gy WT mice. Lastly, to evaluate the impact of cranial RT and C5aR1 inhibition on the hippocampal-amygdala fear memory consolidation circuit, mice were administered a fear extinction test reliant on hippocampal function (21,22). During the conditioning phase (three tones and shocks), all groups of mice showed comparable freezing, indicating the acquisition of conditioned fear memory (**Suppl. Fig. S3**). Interestingly, during the extinction training phase, C5aR1-KO mice (0 and 9 Gy) groups show elevated freezing compared to the WT + 0 Gy + Vehicle group. Such elevated freezing behavior was also observed previously in complement knockout mice by us (microglia-selective C1q knockdown) (3) and others (CR3-KO) (20). A CD55-mediated inhibition of classical complement cascade has suggested the dependence of dissociative learning on the microglial complement signaling (23). We also found significantly increased freezing behavior by the irradiated WT mice (WT + 9 Gy + Vehicle) compared to the WT + 0 Gy controls during the extinction training phase. In the end, during the extinction test phase (**Fig. 1E**), WT + 9 Gy + Vehicle mice showed the highest freezing, an indicator of disrupted memory consolidation processes, compared to WT + 0 Gy + Vehicle mice (P<0.001). The freezing behavior of WT + 0 Gy + PMX205 and C5aR1-KO + 0 Gy mice was comparable to the WT + 0 Gy + Vehicle group. WT + 9 Gy + PMX205 and C5aR1-KO + 9 Gy mice showed reduced freezing compared to WT + 9 Gy + Vehicle (P’s<0.02, **Fig. 1E**), indicating a beneficial cognitive effect. In summary, both genetic knockout and pharmacologic inhibition of the C5a-C5aR1 axis protect against RICD.

### C5aR1 inhibition reduced neuroinflammation

To determine if C5aR1 inhibition via (PMX205 in the WT mice or C5aR1 knockout prevented radiation-induced neuroinflammation, surrogate markers of RT-induced gliosis and complement anaphylatoxins were evaluated using immunofluorescence staining and 3D algorithm-based volumetric quantification of immunoreactivity. Cranial RT significantly elevated microglial lysosomal phagocytosis marker CD68 (CD68-IBA1, **Fig. 2A-C**) in the WT + 9 Gy + Vehicle brains compared to WT + 0 Gy + Vehicle group (P<0.0001). WT + 0 Gy and 9 Gy mice receiving PMX205 treatment showed significantly lower CD68-IBA1 compared to the WT + 9 Gy + Vehicle group (P’s<0.0001, **Fig. 2C**). Similarly, irradiated C5aR1-KO mice (C5aR1-KO + 9 Gy) showed reduced microglial activation compared to WT + 9 Gy + Vehicle group (P<0.0001). Microglial C5aR1 expression was significantly elevated in the brains of WT + 9 Gy + Vehicle compared to WT + 0 Gy + Vehicle, and WT + 0 Gy + PMX205 groups (P’s≤<0.01, **Fig. 2D-F**). PMX205 treatment to the 9 Gy irradiated animals significantly reduced microglial-C5aR1 expression compared to irradiated WT mice receiving vehicle (P<0.001). Conversely, 0 and 9 Gy irradiated C5aR1-KO brains showed minimal C5aR1 immunoreactivity. Cranial RT also significantly elevated astrocytic hypertrophy with thicker and longer GFAP^+^ stelae (green, **Fig. 2G-H**) in Wt + 9 Gy + Vehicle brains compared to WT + 0 Gy + Vehicle group (P<0.01). The GFAP immunoreactivity did not differ between the WT + 0 Gy + Vehicle and WT + 0 Gy + PMX205 groups. PMX205 treatment to the irradiated WT mice (WT + 9 Gy + PMX205) significantly reduced GFAP immunoreactivity compared to WT + 9 Gy + Vehicle group (P<0.002). 0 and 9 Gy irradiated C5aR1-KO mice did not show elevated astrocytic hypertrophy (**Fig. 2G-H**). Thus, C5aR1 inhibition or knockout prevented cranial RT-induced microglial activation and astrogliosis that was coincident with neurocognitive improvements.

**Figure 2.**
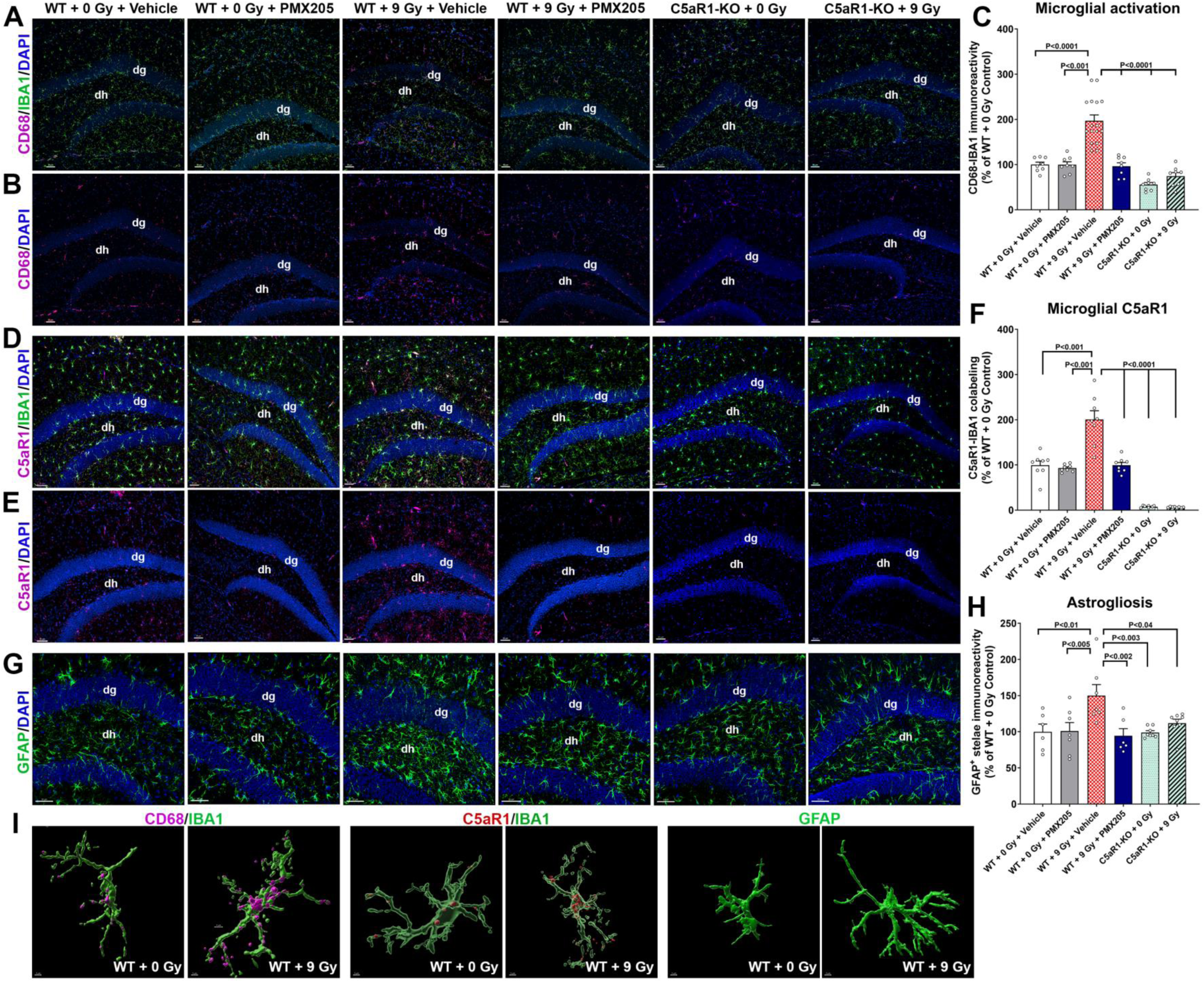
C5aR1 signaling blockade prevents radiation-induced neuroinflammation. **(A)** Dual immunofluorescence staining, confocal microscopy, and 3D algorithm-based, volumetric quantification for the microglial activation (CD68-IBA1 immunoreactivity, magenta and green respectively) in the WT mice show cranial irradiation (WT + 9 Gy + Vehicle)-induced elevated CD68 expression by the IBA1^+^ microglia in the hippocampus (*dh*, dentate hilus; *dg*, dentate gyrus) compared to WT + 0 Gy + Vehicle. **(B)** A separate CD68-DAPI channel (magenta-blue) is shown for each group. **(C)** C5aR1 inhibition (WT + 9 Gy + PMX205) or knockdown (C5aR1 + 9 Gy) did not elevate CD68 immunoreactivity in the irradiated brain, indicating prevention of neuroinflammation. **(D-E)** Volumetric quantification for the microglial C5aR1 expression (C5aR1-IBA1 immunoreactivity, magenta and green, respectively) showed a radiation-induced increase in the microglial-C5aR1 in the WT hippocampal *dg* and *dh* (WT + 9 Gy + Vehicle) compared to unirradiated controls (WT + 0 Gy + Vehicle). C5aR1 inhibition via PMX205 treatment (WT + 9 Gy + PMX205) or the genetic knockout (C5aR1 + 9 Gy) reduced microglial-C5aR1 expression compared to the irradiated WT mice (WT + 9 Gy +vehicle) (**F**). **(G)** Analysis of astrocytic hypertrophy (GFAP^+^ stelae, green) showed radiation-induced elevation in GFAP^+^ astrocytic processes in the WT + 9 Gy+ Vehicle mice compared to WT + 0 Gy + Vehicle or WT + 0 Gy+ PMX205 groups in the hippocampal *dg*. Irradiated WT mice receiving the inhibitor treatment (PMX205) and the irradiated C5aR1-KO mice did not show elevated GFAP immunoreactivity, indicating reduced astrogliosis **(H)**. **(I)** Representative high-resolution 3D volumetric surface rendering derived from immunoreactivity for the activated microglia (CD68-IBA1, magenta-green), microglial C5aR1 expression (C5aR1-IBA1, red-green), and hypertrophic astrocytes (GFAP, green) showing elevated volumes in the WT + 9 Gy + Vehicle group compared to the WT + 0 Gy + Vehicle group. Mean ± SEM (*N*=6-8 per group). Scale bars, 50 µm (**A-B, D-E, G**) and 5 µm (**I**).

### C5aR1 inhibition prevented RT-induced disruption of synaptic integrity

Microglial and complement cascade activation have been shown to play detrimental roles in excessive synaptic loss in neurodegenerative conditions, including RICD (3,6). To determine the impact of C5aR1 blockade using PMX205 or genetic knockout on the synaptic integrity in the irradiated brain, we quantified immunoreactivity of post- and pre-synaptic markers PSD95 and synaptophysin (Syn), respectively (**Fig. 3**). Previously, we reported that RT elevates PSD95 in the mice brains that were cognitively impaired (24,25). Likewise, in this study, we found elevated PSD95^+^ in the irradiated WT hippocampal molecular layer (WT + 9 Gy + Vehicle) compared to WT + 0 Gy + Vehicle and WT + 0 Gy + PMX205 groups (P’s<0.0001, **Fig. 3A-B**). PMX205 treatment to the irradiated WT mice (WT + 9 Gy+PMX205) and C5aR1-KO + 9 Gy mice showed a significant PSD95 reduction compared to WT + 9 Gy + Vehicle mouse brains (P<0.0001). In contrast, RT significantly reduced Syn^+^ puncta in the WT + 9 Gy + Vehicle group compared to the WT + 0 Gy + Vehicle (P<0.0001, **Fig. 3C-D**). The Syn levels were comparable between the WT + 0 Gy + Vehicle and WT + 9 Gy + PMX205-treated mice. Irradiated WT mice receiving PMX205 (WT + 9 Gy + PMX205, P<0.05) and irradiated C5aR1-KO mice (C5aR1-KO + 9 Gy, P<0.0001) did not show reduced Syn compared to the WT + 9 Gy + Vehicle group. Overall, these data indicate a protective role of C5aR1 inhibition against RT-induced disruption of synaptic integrity.

**FIGURE 3.**
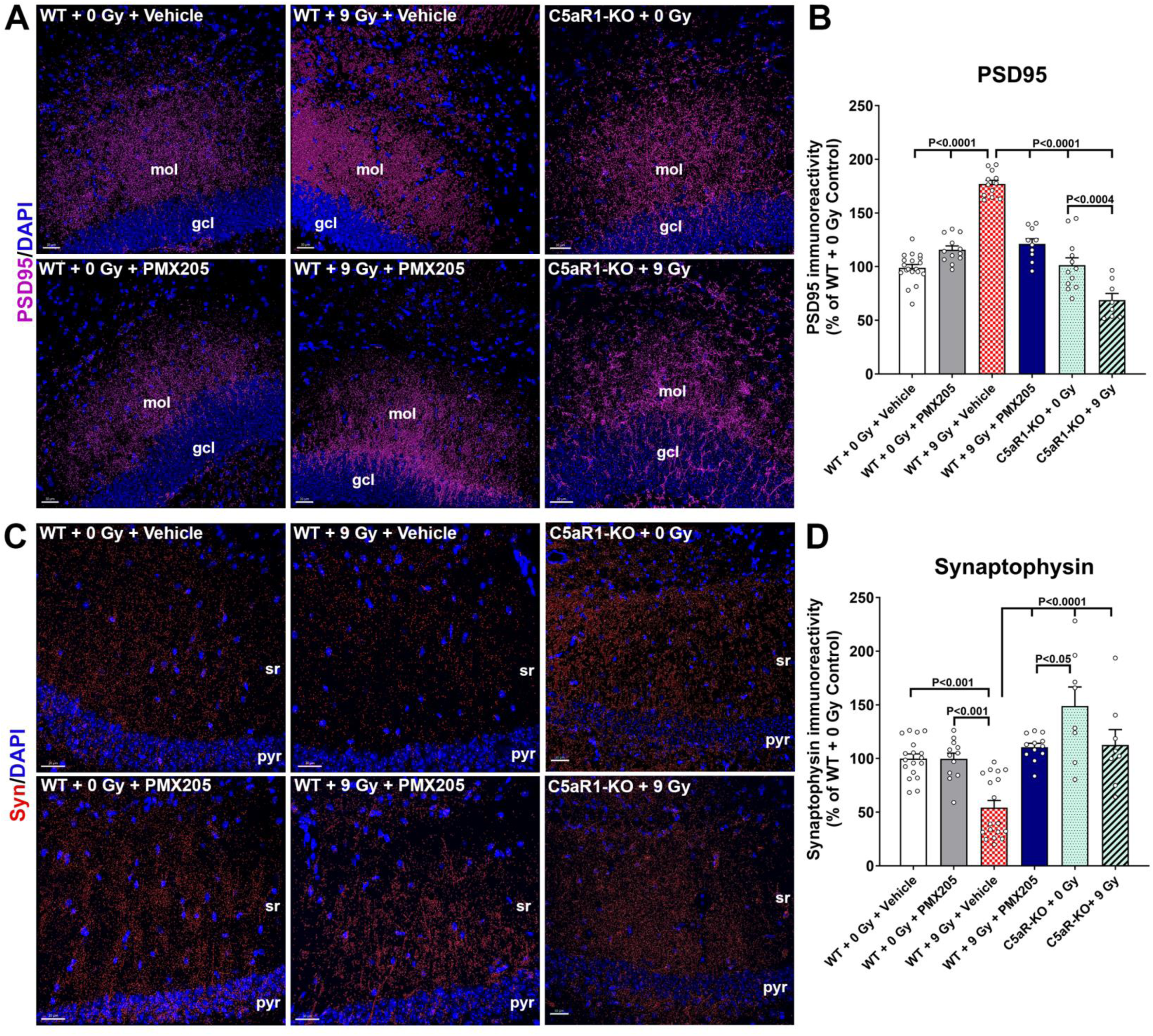
C5aR1 antagonism prevents radiation-induced loss of synaptic integrity. (A-B) Immunofluorescence staining, confocal microscopy, and 3D algorithm-based volumetric quantification for the post-synaptic density protein, PSD95 (magenta, DAPI nuclear counter stain) in the hippocampus (*gcl*, granule cell layer; *mol*, molecular layer) showed significantly elevated PSD95 in WT + 9 Gy+ Vehicle compared to WT + 0 Gy + Vehicle, WT + 0 Gy + PMX205, and C5aR1-KO + 0 Gy groups. Irradiated WT mice receiving a C5aR antagonist (WT + 9 Gy + PMX205) and irradiated C5aR1-KO mice (C5aR1-KO + 9 Gy) did not show an aberrant PSD95 elevation. **(C-D)** In contrast, volumetric quantification of hippocampal synaptic vesicle protein synaptophysin (Syn, red, DAPI nuclear counter stain) showed a significant decline in the hippocampal CA1 (*pyr*, pyramidal layer; *sr*, stratum radiatum) post-RT in the WT mice (WT + 9 Gy + Vehicle) compared to WT + 0 Gy + Vehicle group. PMX205 treatment in the irradiated WT mice (WT + 9 Gy + PMX205) and C5aR1-KO in the irradiated mice (C5aR1-KO + 9 Gy) prevented loss of synaptophysin, indicating the neuroprotective impact of C5aR1 signaling inhibition against cranial irradiation. Mean ± SEM (*N*=8-10 mice per group). Scale bars, 30 µm (**A, C**).

### PMX205 treatment prevented RICD in the brain of cancer-bearing mice without interfering with the therapeutic efficacy of RT

Cranial RT is the primary treatment modality to stall brain cancer progression. The impact of C5aR1 signaling inhibition on WT and C5aR1-KO brain cancer growth, overall survival, and cognitive function post-RT was tested using two orthotopic glioblastoma multiforme (GBM) models. WT mice received intra-cranial injection of syngeneic murine Gl261 or CT2A-luc^+^ cells to model glioblastoma and astrocytoma, respectively (Research design, **Fig. 4A**). One-week post-tumor induction and detection via imaging, mice received 9 Gy cranial-RT. As described previously, one cohort of mice also received PMX205 treatment (1 mg/kg SQ for one week concurrently with 20 μg/ml in the drinking water). Concurrently, C5aR1-KO mice were implanted with CT2A glioma cells and irradiated (9 Gy cranial) 1-week later. For 4-5 weeks post-tumor induction, tumor growth, tumor burden, and animal health status were monitored, followed by the object recognition memory (ORM) behavior test. Gl261 implanted mice showed a steady tumor growth as detected by the iodine contrast (omnipaque) CBCT (**Fig. 4B**). Compared to WT + Gl261 + Vehicle group, the tumor growth in the WT + Gl261 + 9 Gy and WT + Gl261 +PMX205 (black line) was significantly lower at week 3-5 post-induction (P’s≤0.05). Conversely, mice implanted with CT2A-luc showed a rapid and steady tumor progression in the 0 and 0 Gy cranially irradiated WT brains as detected by the bioluminescence imaging (BLI, **Fig. 4C**). Irradiated C5aR1-KO mice with tumor (C5aR1-KO + CT2A + 9 Gy) showed significantly reduced tumor growth compared to WT + CT2A + Vehicle mice (P’s≤0.05, **Fig. 4C**). At 4-weeks post-implantation, we found a significant reduction in the tumor burden in the WT mice receiving 9 Gy RT and PMX205 treatments (WT + CT2A+ 9 Gy, WT + CT2A + PMX205, and WT + CT2A + 9 Gy + PMX205) compared to the CT2A+Vehicle group (P’s<0.001).

**FIGURE 4.**
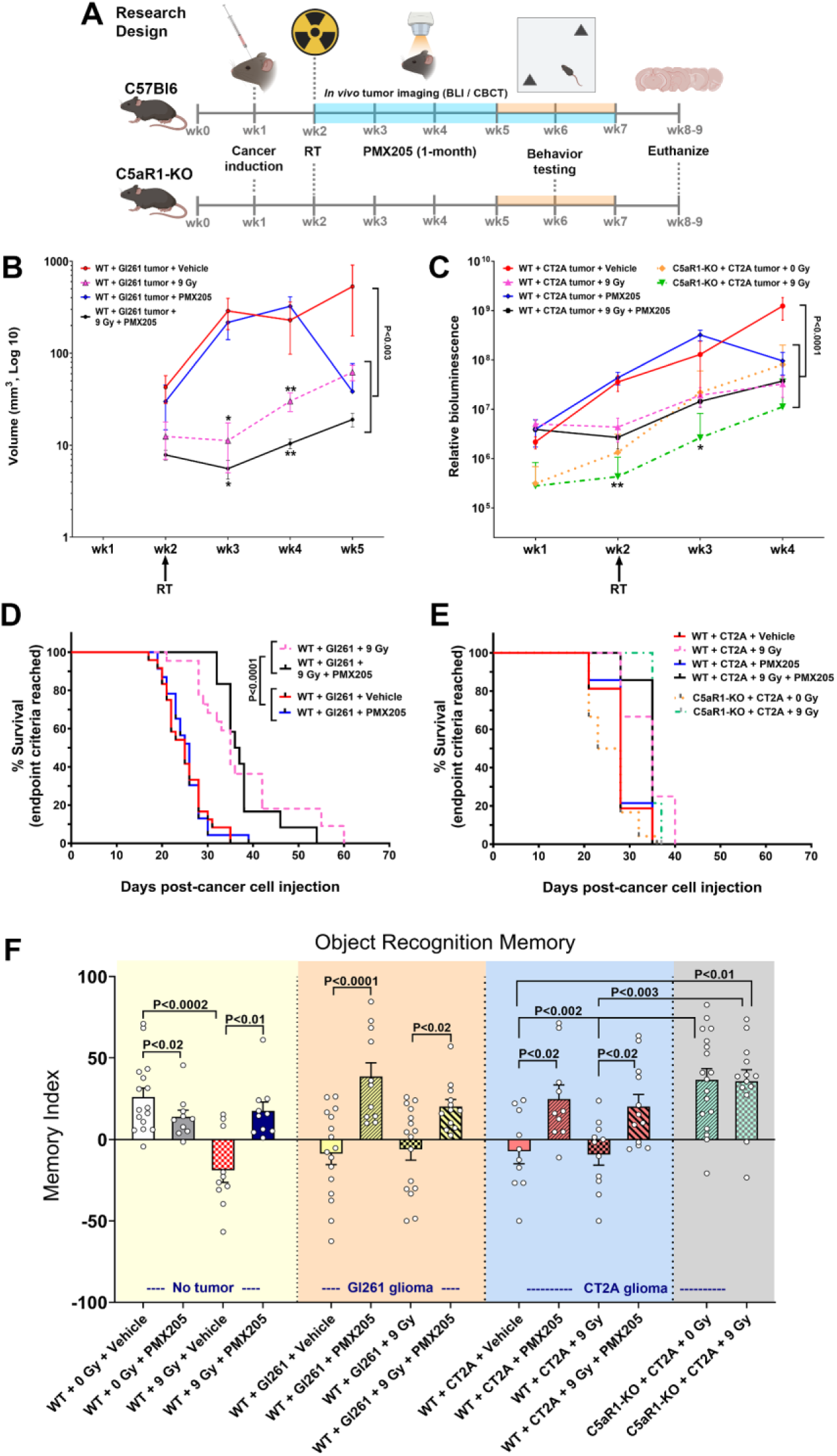
C5aR1 signaling inhibition in the glioma-bearing mice reversed cranial radiation-induced cognitive decline without interfering with the therapeutic efficacy of RT. **(A)** Schematic representation of research design. C57Bl6 mice received orthotopic implantation of syngeneic cancer cells (Gl261, and CT2A-luc) in the caudate putamen. A cohort of brain cancer-bearing WT mice received cranial radiotherapy (RT, 9 Gy) 1-week post-tumor induction. Mice were divided into the following groups (2 tumor cell types): Tumor + Vehicle, Tumor + RT, Tumor + PMX205, and Tumor + RT + PMX205. At 48h post-RT, mice received PMX205 treatment (1 mg/kg SQ for the first week in addition to 20 μg/ml, or 100 μg per day in drinking water for 1-month). A separate cohort of transgenic C5aR1-KO mice received CT2A-luc cell implantation ± RT. Tumor growth and progression were monitored weekly by *in vivo* tumor imaging. Bioluminescence Imaging (BLI) was used for CT2A-luc^+^ glioma, and iodine contrast (Omnipaque), Cone-Beam Computerized Tomography (CBCT) scanning was used for the detection and volumetric measurement of Gl261^+^ glioblastoma. At 3 to 4 weeks post-RT, mice were administered object recognition memory task (ORM) and euthanized for tissue collection. **(B)** CBCT-acquired Gl261^+^ glioblastoma showed a steady progression through one-week post-implantation and RT. Gl261 glioma-bearing mice receiving RT (WT + Gl261 + 9 Gy) and WT + Gl261 + 9 Gy + PMX205 groups showed a significant reduction in tumor growth post-RT compared to WT + Gl261 + Vehicle and WT + Gl261 + PMX205 groups. At the end of the study (5 weeks post-tumor implantation), the WT + Gl261 + Vehicle group showed a significantly higher bioluminescence signal than all the other WT groups. *P<0.05, and **P<0.01 compared to WT + Tumor + Vehicle group. **(C)** The CT2A-luc^+^ astrocytoma showed rapid growth 1-week post-implantation. C5aR1 inhibition (PMX205) in the WT mice (CT2A + PMX205, CT2A + 9 Gy, and CT2A + 9 Gy + PMX205), and C5aR1-KO mice receiving RT showed a significant reduction in tumor volume compared to the CT2A + vehicle group. *P<0.05, and **P<0.01 compared to WT + CT2a + Vehicle group. **(D-E)** Kaplan-Meier estimates showed improved survival of Gl261^+^ glioblastoma-bearing mice receiving RT or 9 Gy + PMX205 treatments compared to Vehicle or PMX205 treatment alone. CT2A^+^ astrocytoma showed comparable survival estimates for all groups of mice. See **Suppl. Fig. S3** for C5aR-KO and WT group survival. (F) At 3-4 weeks post-RT, cognitive function assessment (ORM task) was performed. To compare cognitive function, intact (no tumor) WT mice performance (as in Fig. 1C) is plotted along with glioma groups. 9 Gy cranial RT significantly reduced memory index (WT + 9 Gy + Vehicle) in the WT mice without tumors. Tumor burden itself (Gl261+Veh and CT2A+Veh) and tumor-bearing mice receiving RT (WT + Gl261+ 9 Gy and WT + CT2A + 9 Gy) showed a significant reduction in the ORM task performance (reduced Memory Index). In contrast, tumor-bearing WT mice ± 9 Gy receiving C5aR1 inhibitor (PMX205) treatment and CT2A-bearing C5aR1-KO mice ± 9 Gy showed significant improvements in Memory Indices (WT + Gl261 + PMX205, WT + Gl261 + 9 Gy + PMX205, WT + CT2A + PMX205, WT + CT2A + 9 Gy + PMX205, C5aR1-KO + 0 Gy, and C5aR1-KO + 9 Gy), indicating neuroprotective impact of C5aR1 signaling blockage. Mean ± SEM (*N*=12-24 mice per group).

The Kaplan-Meier survival curves for the Gl261-induced glioma model showed improved survival rates for the GBM-bearing WT mice receiving 9 Gy RT or RT + PMX205 treatments (WT + Gl261 + 9 Gy, and WT + Gl261 + 9 Gy + PMX205) compared to WT GBM mice treated with vehicle or PMX205 without RT (P’s<0.0001, **Fig. 4D**). On an average, survival for Gl261-bearing WT mice receiving RT or RT + PMX205 treatments was 54-60 days, whereas, vehicle or PMX205 treatment alone did not influence the survival beyond 36-39 days post-tumor induction. Contrary to this, the CT2A-luc-implanted WT and C5aR1-KO glioma model did not show significant improvements in survival following 9 Gy RT alone, PMX205 alone, or 9 Gy RT + PMX205 treatments, and the average survival was between 35-40 days post-tumor induction (**Fig. 4E**). A separate survival curve comparing C5aR1-KO transgenic and WT backgrounds for CT2A-induced glioma ± RT (derived from **Fig. 4E**) is shown in **Suppl. Fig. S4**.

Concurrently, at 3-4 weeks post-RT, neurocognitive function of the brain cancer-bearing WT mice with or without PMX205 treatment was assessed using the ORM task (**Fig. 4F**). For reference, ORM data for *No Tumor* mice was derived from **Fig. 1C**. Tumor burden significantly reduced the performance on the ORM task for both WT + Gl261 + Vehicle and WT + CT2A + Vehicle groups compared to WT + 0 Gy + Vehicle (P’s≤0.01, **Fig. 4F**, *Gl261 glioma* and *CT2A glioma* groups). Cancer-bearing WT mice receiving RT (WT + Gl261 + 9 Gy and WT + CT2A + 9 Gy) also performed poorly on the ORM task (P’s≤0.01) versus WT + 0 Gy + Vehicle. Importantly. Gl261- and CT2A-bearing WT mice treated with PMX205 showed significantly improved memory index compared to Gl261- and CT2A-bearing WT mice + Vehicle (P<0.0001, P<0.02 respectively, **Fig. 4F**). Remarkably, PMX205 treatment significantly improved memory indices for the brain cancer-bearing WT mice receiving RT. The performance of WT + Gl261 + 9 Gy + PMX205 and WT + CT2A + 9 Gy + PMX205 mice was significantly higher compared to WT + Gl261 + 9 Gy and WT + CT2A + 9 Gy groups respectively (P’s<0.02). This data was further corroborated by intact object recognition memory function by the C5aR1-KO + CT2A + 0 Gy and C5aR1-KO + CT2A + 9 Gy groups compared to WT + CT2A + Vehicle and WT + CT2A + 9 Gy groups. (**Fig. 4F**). Thus, PMX205 treatment and C5aR1 knockout were neuroprotective in brain cancer-bearing mice receiving cranial RT.

### C5aR1 inhibition or knockout reduced radiation-induced glial activation and loss of synaptic integrity in the glioma-bearing brains

Neurocognitive function data using pharmacologic and genetic approaches indicated the beneficial impact of ablating C5aR1 signaling in the brains of glioma-bearing, irradiated mice, thus alleviating normal tissue toxicity impact of cranial RT (**Fig. 4**). Moreover, irradiated WT and C5aR1-KO brains without tumor were protected from RT-induced neuroinflammation, anaphylatoxin receptor expression and the loss of synaptic integrity (**Figs. 2-3**). To determine if C5aR1 blockage was equally neuroprotective in the brains of CT2A-induced glioma-bearing mice, we assessed microglial activation (CD68-IBA1, **Fig. 5A-C**), microglial C5aR1 expression (C5aR1-IBA1, **Fig. 5D-F**), and astrogliosis (**Fig. 5G-H**) in the irradiated WT and C5aR1-KO hippocampus. Glioma burden significantly elevated the volume of microglial CD68 (**Fig. 5A-C**), C5aR1 immunoreactivity (**Fig. 5D-F**), and astrocytic hypertrophy with thicker and longer GFAP^+^ stelae (**Fig. 5G-H**) in the WT + CT2A + Vehicle and WT + CT2A + PMX205 groups compared to WT + 0 Gy + Vehicle group (P’s≤0.01). Conversely, WT + CT2A + 9 Gy mice treated with PMX205 and C5aR1-KO + CT2A mice receiving either 0 or 9 Gy cranial RT showed significantly reduced gliosis or the C5aR1 expression (P≤0.01) indicating reduced neuroinflammation. A multiplex ELISA-based estimation for a cytokine panel from hippocampal extracts did not reveal significant differences between the experimental groups except for IFNᵧ, which was found to be significantly elevated in the WT + CT2A + Vehicle and WT + CT2A + 9 Gy hippocampus (**Suppl. Fig. S5**). We also found reduced hippocampal synaptophysin (Syn) expression in the WT + CT2A + Vehicle, WT + CT2A + 9 Gy, and WT + CT2A + PMX205 groups compared to WT + 0 Gy + Vehicle (**Fig. 5I-J**, P’s≤0.01). The oral treatment with PMX205 to the WT + CT2A + 9 Gy mice and irradiated C5aR1-KO + CT2A mice showed restored Syn immunoreactivity (P’s≤0.01). On the other hand, PSD95 levels were significantly elevated in the WT + CT2A + 9 Gy hippocampus compared to the WT + 0 Gy + Vehicle group (P<0.0001, Fig. **5K-L**). PMX205 treatment to the WT + CT2A + 9 Gy group and 0 or 9 Gy irradiated C5aR1-KO + CT2A group showed reduced PSD95 (P≤0.01), indicating neuroprotective effects of ablating C5aR1 signaling in the glioma-bearing brains against RT-induced loss of synaptic integrity.

**FIGURE 5.**
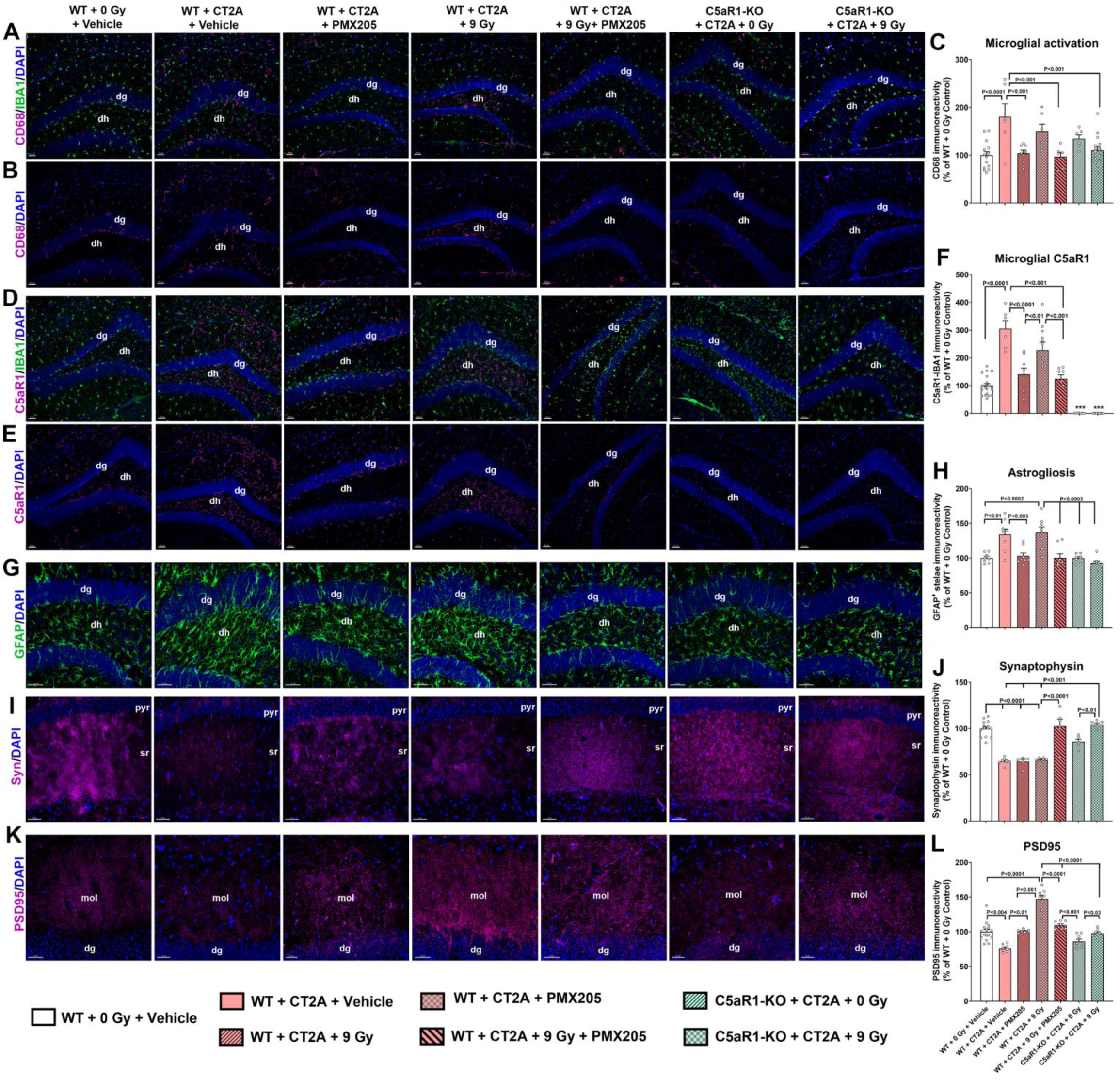
C5aR1 antagonism in the glioma-bearing brain prevents radiation-induced inflammation and the loss of synaptic integrity. **(A-F)** Dual immunofluorescence staining, confocal microscopy, and 3D algorithm-based, volumetric quantification for the microglial activation (CD68, magenta and IBA1 green, **A-B**) and microglial C5aR1 expression (C5aR1, magenta, and IBA1, green, **D-E**) in the CT2A glioma-bearing WT brain showed RT-induced elevated CD68 and C5aR1 expression in the WT + CT2A + Vehicle hippocampus (*dh*, dentate hilus; *dg*, dentate gyrus) compared to WT + 0 Gy + Vehicle. **(B** and **E)** A separate CD68-DAPI channel (magenta-blue) is shown for each group. **(C** and **F)** C5aR1 inhibition (WT + CT2A + 9 Gy + PMX205) or knockdown (C5aR1-KO + CT2A + 9 Gy) did not elevate CD68 and C5aR1 immunoreactivity in the irradiated brain, indicating reduced inflammation. **(G-H)** Similarly, volumetric quantification of astrocytic hypertrophy (GFAP^+^ stelae, green) showed elevated GFAP^+^ astrocytic processes in the WT + CT2A + Vehicle and WT + CT2A + 9 Gy + Vehicle mice compared to WT + 0 Gy + Vehicle or WT + 0 Gy+ PMX205 groups in the hippocampal *dg*. Irradiated WT mice receiving the inhibitor treatment (WT + CT2A + 9 Gy + PMX205) and the irradiated C5aR1-KO mice (C5aR1-KO + CT2A + 9 Gy) did not show increased GFAP immunoreactivity, indicating reduced astrogliosis **(H)**. **(I-J)** Volumetric quantification of the hippocampal synaptophysin (syn, magenta, DAPI nuclear counter stain) showed a reduced immunoreactivity in WT + CT2A ± PMX205 and WT + CT2A + 9 Gy groups compared to WT + 0 Gy + Vehicle groups in the hippocampal CA1 pyramidal (*pyr*) and stratum radiatum (*sr*) regions. C5aR1 inhibition in the WT brain (WT + CT2A + 9 Gy + PMX205) and C5aR1 knockout (C5aR1-KO + CT2A + 0 Gy and C5aR1-KO + CT2A + 9 Gy) significantly increased synaptophysin **(J)**. **(K-L)** On the other hand, volumetric quantification of the hippocampal PSD95 (magenta, DAPI nuclear counterstain) showed radiation-induced elevated immunoreactivity (*mol*, molecular layer; *dg*, dentate gyrus) in the WT + CT2A + 9 Gy group compared to WT + 0 Gy + Vehicle. Irradiated WT mice receiving a C5aR1 antagonist (WT + CT2A + 9 Gy + PMX205) and irradiated C5aR1-KO mice (C5aR1-KO + CT2A + 9 Gy) showed reduced PSD95. (I-L) The synaptic protein expression data indicates intact synaptic integrity in the cancer-bearing irradiated brains following C5aR1 antagonism. Mean ± SEM (*N*=6-16 per group). Scale bars, 50 µm (**A-E,** and **K**), 40 µm (**G and I**).

### C5aR1 signaling blockade promotes neuroprotective gene expression in the irradiated brain

The preceding data showed the neuroprotective role of C5aR1 inhibition or knockout in the intact non-cancer and cancer-bearing mouse brain receiving cranial RT (**Figs. 1** and **4**). The brain function outcomes were also supported by reduced glial activation and loss of synaptic integrity (**Figs. 2-3** and **5**). To further corroborate these findings, we investigated the expression of neuroinflammation-related genes using a NanoString neuroinflammation panel with a set of 757 genes. RNAs were extracted from WT, WT + PMX205 and C5aR1-KO mice brains with or without 9 Gy cranial RT and with or without CT2A-induced gliomas after the completion of behavior studies. This analysis aimed to determine if PMX205 treatment or C5aR1 knockout reduces the neuroinflammatory environment in the irradiated brains and thereby preventing RT-induced normal tissue toxicity. We compared WT + 9 Gy + PMX205 and C5aR1-KO + 9 Gy with the WT + 9 Gy + Vehicle group as a baseline separately without induction of tumors (**Fig. 6**). The adjusted P≤0.05 was the cutoff criteria. Amongst 55 differentially expressed genes (DEGs), we found 13 upregulated and 33 downregulated genes in the WT + 9 Gy + PMX205 group compared to the WT + 9 Gy + Vehicle group (**Fig. 6A-D**, and **Suppl. Table T1**). In contrast, out of 60 DEGs, 46 genes were upregulated, and 5 genes were downregulated in the C5aR1-KO + 9 Gy group compared to WT + 9 Gy + Vehicle. The 9 DEGs were shared between the WT + 9 Gy + PMX205 and C5aR1-KO + 9 Gy brains (**Fig. 6E-F**, and **Suppl. Table T1**). *Rps10*, a ribosome component, was downregulated in the brains of WT + 9 Gy + PMX205 and upregulated in C5aR1-KO + 9 Gy mice. The shared DEGs (**Fig. 6F**) were affiliated with microglial function, neurotransmission, and DNA damage response (**Suppl. Table T1**). Comparison of WT + 9 Gy + Vehicle and WT + 0 Gy + Vehicle found upregulation of *Chn2 in vivo* (P<0.05, **Supplemental Fig. S6 A**). Whereas comparison of CT2A glioma-bearing WT and C5aR1-KO brains receiving 9 Gy RT with WT + CT2A + 9 Gy did not reveal statistical significance for the DEGs (**Supplemental Fig. S6 B-C**).

**FIGURE 6.**
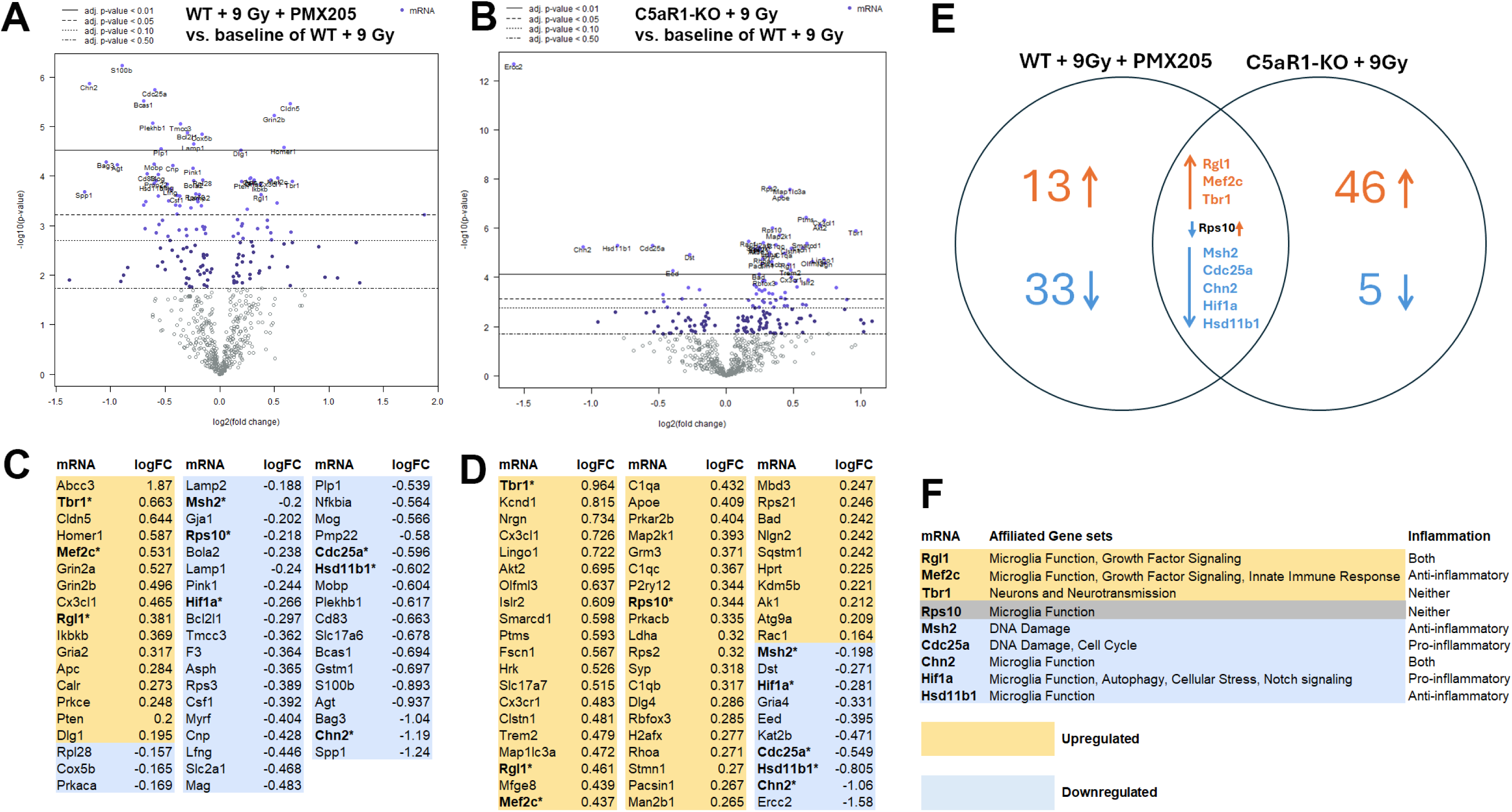
C5aR1 knockout or inhibition downregulates neuroinflammatory gene expression in the irradiated brain. **(A-B)**: Volcano plots of differentially expressed genes (DEGs) comparing WT + 9 Gy + PMX205 (**A**) and C5aR1-KO + 9Gy (**B**) to the WT + 9 Gy + Vehicle group as a baseline. Brain RNA was analyzed 8-9 weeks post-irradiation using the NanoString nCounter Neuroinflammation panel. The log2 fold change (logFC) is plotted against the -log10 (adjusted P value) for each gene that met the minimal count threshold. Amongst 757 genes, 745 met count thresholds in all animal groups. **(C-D)**: The list of topmost significant DEGs for both WT + 9 Gy + PMX205 (**C**) and C5aR1-KO + 9Gy (**D**) groups showing significant differences (adjusted P<0.05). The bold-faced (**\***) genes were shared between both treatment groups and showed comparable differential expression in the irradiated brains. Yellow highlight upregulated (logFC>0) and blue were downregulated (logFC<0) genes. C5aR1-KO + 9Gy DEGs showed a larger proportion of upregulation compared to WT + 9Gy + PMX205. See **Supplemental Table T1** for gene function details. **(E)** Venn diagram showing the number of DEGs (adjusted P<0.05) for the two treatment groups: WT + 9 Gy + PMX205 and C5aR1-KO + 9Gy. Amongst 55 DEGs, WT + 9 Gy + PMX205 had 46 unique, 13 upregulated, and 33 downregulated DEGs compared to the WT + 9 Gy + Vehicle group. On the other hand, out of 60 DEGs, C5aR1-KO + 9Gy brains had 51 unique, 46 upregulated, and 5 downregulated genes, showing a greater impact of genetic knockdown of the C5aR1 in the irradiated tissue bed. **(F)** Shared DEGs between WT + 9 Gy + PMX205 and C5aR1-KO + 9Gy treatment groups (WT + 9 Gy + Vehicle group as a baseline) and their affiliated gene sets as analyzed by the NanoString nCounter Advanced Analysis (criteria: adjusted P<0.05). C5aR1 signaling ablation upregulates anti-neuroinflammatory and neuroprotective genes and downregulates inflammation, cell, and genomic instability-related genes (**Suppl. Table T1**). Adjusted P values (P≤0.05) were derived from the Benjamini-Yekutieli False Discovery Rate (FDR) method (N = 4 brains per group).

## DISCUSSION

RICD, prevalent in 50-90% of surviving brain cancer patients, has long been recognized as adversely afflicting their overall quality of life (QOL)(26). Yet, neither neurobiological mechanisms driving such poor neurocognitive outcomes nor preventative measures are well developed. Despite extensive knowledge of aberrant complement activation in neurodegenerative conditions, specific consequences of the C5a-C5aR1 signaling axis in the irradiated and cancer-bearing brain are not yet reported. RT-induced microglial activation plays a pivotal damaging role in leading to a cascade of pro-inflammatory events, including CNS complement activation (3,6). The current study sought to understand the detrimental role of the downstream complement C5a signaling. C5a is an anaphylatoxin and a product of complement cascade activation via cleavage of C5. It has been shown to stimulate chemotaxis and a robust pro-inflammatory response after interaction with C5aR1 (6). Activation of the C5a-C5aR1 axis promotes pro-inflammatory polarization of microglia, excessive synaptic pruning, and loss of cognitive function in AD models (7,9). C5a overexpression in AD mice led to accelerated hippocampal-dependent memory loss (10). Contrarily, genetic KO or selective C5aR1 inhibition protected the AD brain against microglia-mediated excessive synaptic loss, improved neuronal function, and reduced dementia (7,9,27). These reports established the detrimental role of the C5a-C5aR1 axis in promoting cognitive decline in a chronic neurodegenerative condition. Cranial RT led to a similar extent of microglial activation, synaptic loss, and cognitive decline (sans plaque deposition) in the brain (3,24,28,29). Cranial RT-induced cognitive impairments were linked with elevated microglial activation, astrocytic hypertrophy, and elevated pro-inflammatory cytokines *in vivo* (28,30,31). Subsequently, microglia elimination or reduced astrogliosis reversed those cognitive deficits (28,32). CNS complement components are co-localized or expressed by both microglia and astrocytes *in vivo*, and they synchronize pathologic response and normal tissue toxicity in the irradiated brain.

We first sought to understand an overarching issue of an RT-induced neuropathologic condition in intact, cancer-free mouse brains. We aimed to determine if classical complement cascade activation and formation of anaphylatoxin steers microglial and astrocytic activation, leading to excessive synaptic alterations and cognitive decline. To prove the principle, we tested two approaches, genetic C5aR1 KO and pharmacologic inhibition, using a selective, orally active inhibitor, PMX205. Exposure to an acute 9 Gy cranial RT led to a significant disruption in hippocampal, frontal cortex, and amygdala-dependent learning and memory and memory consolidation and re-learning processes (**Fig. 1**). In the past, we have shown that such neurocognitive disruptions are often progressive, persistent for as long as 8-months post-RT and do not resolve over a time (30,33). RICD was also reflected in the results of the fear extinction memory test that measures memory consolidation, an ability to re-learn or dissociate from a previously aversive experience. Importantly, PMX205 significantly reduced RICD. Comparable results in the C5aR1 KO mice asserted the selective specificity of our pharmacologic C5aR1 inhibition approach. Irradiated C5aR1 KO mice did not show a cognitive decline compared to the irradiated WT mice. These data support our previous findings that show accelerated cognitive impairments following C5a overexpression (10) and that C5aR1 inhibition (PMX205) improved cognition in AD model (7). Among the recent advances targeting complement activation post-RT, our work has shown that microglia-selective deletion of C1qa in the irradiated CNS prevented RICD and excessive synaptic loss(3). Moreover, inhibition of complement receptor 3 (CR3) axis also reduced RICD (20) in male mice. Such inhibition of upstream complement C1q and C3 would prevent downstream events such as C5a generation C5aR1 signaling (3,6). As these upstream complement components play several beneficial roles, a complete ablation or inhibition of the CNS complement cascade may not be an optimal mitigation strategy for the irradiated brain. Instead, inhibition of the downstream anaphylatoxin C5a-C5aR1 signaling axis is a better approach to prevent microglial activation and the RICD.

In this study, we confirmed the status of microglial activation, microglial-C5aR1 expression, and astrogliosis by dual-immunofluorescence and 3D algorithm-based volumetric quantification (3) of immunoreactive cell surface or puncta and *in silico* quantification of colocalization post-C5aR1 inhibitor treatment in the WT and C5aR1-KO irradiated brains (**Fig. 2**). Cranial RT-induced elevated microglial activation was characterized by increased expression of the lysosomal phagocytotic marker, CD68, by the IBA^+^ microglia. The microglial C5a receptor (C5aR1-IBA1) expression was also elevated post-RT. The irradiated brains also showed hypertrophic astrocytes with thicker and longer cellular processes. Together, these markers are characteristic of gliosis in the irradiated brain. C5aR1 knockout or treatment with PMX205 in the irradiated brain significantly reduced microglial activation, C5aR1 expression, and astrogliosis. Concurrently, the assessment of synaptic proteins showed two opposite trends. RT significantly elevated PSD95 and reduced synaptophysin (**Fig. 3**), and conversely, irradiated WT brains treated with PMX205 and C5aR1 knockout showed restoration in this synaptic protein expression. These data corroborate our past findings that cranial RT exposure increased PSD95 and reduced synaptophysin expression (24). PSD95 is a scaffolding protein that plays important roles in neuron and dendritic spine growth and maturation, yet mutations and varied post-translational modifications in PSD95 are also linked with neurodevelopmental disorders (34,35). Aging and neuroinflammation-related elevations in PSD95 were correlated with imbalanced excitatory to inhibitory synaptic circuits, leading to excitotoxic events and neuropsychiatric disorders (36-38). Cranial RT-induced elevated PSD95 was suggested to coincide with reduced dendritic branching, dendritic complexity, and reduced expression of presynaptic marker synaptophysin (39). Our data showed restoration in the expression of these synaptic proteins in the irradiated C5aR1-KO and the WT + RT + PMX205-treated brains. C5aR1 signaling is critical in complement-mediated microglial synaptic pruning in the AD brain (9). Furthermore, analyses of Arctic (AD) mice brains either crossed with C5aR1 KO(7) or treated with PMX205 (8) showed a shift in the microglial polarization to a less inflammatory, more phagocytic state that facilitated plaque clearance and reduced gliosis. The C5aR1 signaling axis regulates both microglial functions, complement-mediated synaptic pruning, and synaptic engulfment (7,40). Elevated microglial C5aR1 expression in the irradiated brain also synergizes with TLR4 expression, DAMP response^3^, and astrogliosis that polarizes microglia to an inflammatory and damaging state, leading to imbalanced synaptic function, thereby leading to RICD. Thus, C5a-C5aR1 axis inhibition will suppress the microglial pro-inflammatory polarization and promote beneficial activity, including clearance of dead cell debris. This possibility still needs to be tested in the irradiated brain. In total, our data and the literature support a neuroprotective role upon inhibition of the C5a-C5aR1 axis in alleviating neurodegenerative consequences post-RT.

The translational feasibility of this neuroprotective strategy against RICD and neuroinflammation needed to be tested in brain cancer. Cranial RT is a clinical treatment regimen for primary and metastatic CNS cancers. The detrimental CNS impact of RT, particularly for the long-term survivors of childhood brain cancer and LGG, reduces the QOL and thus limits the therapeutic benefit (1). We tested the neuroprotective efficacy of C5aR1 inhibition on two syngeneic mouse glioma models using CT2A cells (astrocytic origin) and Gl261 cells (glioblastoma, GBM). Brain cancer-bearing mice receiving cranial RT showed dampened tumor growth compared to unirradiated mice (**Fig. 4**). PMX205 treatment of Gl261 glioma-bearing mice ± RT significantly reduced the GBM growth. In parallel, the irradiated C5aR1-KO environment also reduced CT2A^+^ glioma compared to WT + CT2A + Vehicle mice. Syngeneic GL261 is an aggressive form of high-grade glioma that typically responds to high doses of fractionated RT and temozolomide treatments. In contrast, CT2A^+^ glioma showed a rapid initial growth and radiation response following an acute 9 Gy RT, PMX205 treatment, or C5aR1 knockout. Nonetheless, combined cranial RT and PMX205 treatment improved survival in Gl261-bearing mice compared to the unirradiated cohort. We did not observe significant differences in survival for astrocytoma-bearing WT or C5aR1-KO mice. Overall, these brain cancer data suggest that C5aR1 signaling inhibition did not interfere with the therapeutic outcome of acute (9 Gy) cranial RT.

Clinical studies have shown complement deposition (C3, C5b-9) and overexpression of C3aR in patient-derived GBM tissues (41,42). Elevated complement components have been reported in vascularization and malignant glioma growth (43). Genetic ablation of C3 or C5aR1, and treatment with C5aR1 inhibitor (PMX53) reduced tumor growth and angiogenesis in ovarian cancer (15). PMX205 showed anti-cancer activity and increased effectiveness in RT for intestinal cancer (44). Upregulation of C5aR1 expression has been linked with glioma pathogenesis. A recent study evaluating the TCGA and CGGA databases found a significant upregulation of C5aR1 expression in the LGG and GBM tissues (45). C5aR1 expression increased with increasing the grade of glioma and was linked with poor prognosis (low survival rate). Importantly, PMX205 treatment significantly reduced glioma growth in a xenograft glioma model. Using the two syngeneic mouse glioma models, Gl261 and CT2A, our data corroborated these findings, showing significantly reduced tumor volume *in vivo* and improved survival (Gl261) following the C5aR1-KO or combined treatment with PMX205 and RT (**Fig. 4B-E**). Collectively, C5a-C5aR1 axis inhibition has shown promising anti-cancer efficiency and protection of normal tissue. These reports led us to test cognitive function in our murine brain cancer and cranial RT models to determine if C5aR1 knockout or inhibitor treatment alleviated RT-induced normal tissue toxicity. In our study, glioma burden itself reduced cognitive function compared to unirradiated cancer-free controls (**Fig. 4F**). Cranial RT to glioma-bearing mice also impaired cognitive function. Importantly, PMX205 treatment to WT-bearing mice with or without RT and C5aR1-KO in the cancer-bearing mice receiving RT significantly ameliorated RICD. This is a significant result, and it reinforces findings from the cancer-free irradiated mice (**Fig. 1**), which validate the neuroprotective effect of C5aR1 signaling blockage on cognitive function.

To further validate the neuroprotective efficacy of blocking C5aR1 signaling, the transcriptomic profiling of a panel of 757 neuroinflammation-related genes was conducted (**Fig. 6**). PMX205 treatment to the 9 Gy irradiated WT brain led to upregulation of 13 and downregulation of 33 differentially expressed genes (DEGs) compared to the brains of 0 Gy WT control. A more noticeable impact was observed in the irradiated C5aR1-KO brain, with an upregulation of 46 and a downregulation of 5 DEGs compared to the irradiated WT brain. The brains of C5aR1 inhibitor or knockout mice shared 3 upregulated DEGs, including *Mef2c, Rgl1,* and *Tbr1* (**Fig. 6F**). *Mef2c* is a transcription factor that plays roles in developing neurons, synaptic plasticity, maintaining homeostatic function of microglia, and preventing overactivation(46,47). The neuroprotective effects of *Mef2c* have been shown in neurodegenerative AD models(48). *Rgl1* indirectly regulates microglial cytokines release through Ral GTPase and vesicular trafficking, helps resolve inflammation through the secretion of anti-inflammatory factors, and controls neural progenitor maintenance (49). *Tbr1*, a transcription factor, is involved in brain development, particularly in the differentiation of excitatory (glutamatergic) neurons and maintaining neuronal function (50,51). Conversely, 5 downregulated DEGs were shared between WT + 9 Gy + PMX205 and C5aR1-KO + 9 Gy irradiated brains, including *Cdc25a, Chn2, Hif1a, Hsd11b1,* and *Msh2* **(Fig. 6F)**. *Cdc25a* regulates the G1 to S phase transition of the cell cycle and promotes apoptosis and neurodegeneration in AD, ischemic brain injury, and Parkinson’s disease models (52,53). Inhibition of *Cdc25a* expression reduced neuronal death in the AD brain (54). *Hif1a* promotes microglial activation, the release of IL-1β, and the pro-inflammatory polarization of microglia, leading to neuronal damage (55,56). On the other hand, *Hsd11b1* inhibition or knockout protects against hippocampal inflammation from LPS-induced microglial activation and corticosteroid- or aging-associated cognitive decline (57). *Msh2* is involved in DNA mismatch repair and damage response. Radiation-induced DNA damage and genomic instability could also trigger *Msh2* expression *in vivo (58)*. Downregulation of Msh2 indicates reduced genomic damage following either PMX205 treatment or C5aR1 knockout. *Rps10*, a ribosomal component essential for protein synthesis, was found to be upregulated in the irradiated WT brain + PMX205 and downregulated in the brains of C5aR1-KO + 9 Gy, showing differences between the WT and genetic backgrounds. *Chn2* is linked with microglial trafficking, pro-inflammatory activation, and the production of cytokines (55). *Chn2* also mediates axonal pruning and synapse elimination in the hippocampal dentate gyrus (59). Interestingly, a pairwise comparison of WT + 0 Gy and WT + 9 Gy brains showed significantly elevated *Chn2* in the cranially irradiated animals (**Suppl. Fig. S6**). In addition, WT + CT2A + Vehicle and WT + CT2A + 9 Gy brains also had higher pro-inflammatory IFNᵧ *in vivo* (**Suppl. Fig. S5**). Thus, reduced *Chn2* following C5aR1 signaling blockage indicates a neuroprotective impact of this approach. We did not find significant changes in the DEGs in glioma-bearing, irradiated WT, or C5aR1-KO brains compared to the WT + glioma brains (**Suppl. Fig. S6**). This could be attributed to animal-to-animal variations related to *in vivo* tumor growth, whole brain RNA isolation versus region- and cell type-specific changes, radiation response, and C5aR1 blocking paradigms. Overall, C5aR1 inhibitor treatment or knockout in tumor-free mice upregulated neuroprotective and downregulated pro-inflammatory genes *in vivo*.

Our study demonstrates that targeting C5aR1 in the irradiated brain will not disrupt the beneficial effects of the upstream complement components but will prevent microglial activation and astrogliosis and, importantly, protect from cranial RT-induced normal tissue toxicity and RICD in cancer-free and cancer-bearing brains. C5aR1 inhibitors have shown protective effects in many neurodegenerative and inflammatory disease models, including asthma, sepsis, ischemia, Huntington’s disease, ALS, and TBI (60-65). Human studies using an anti-C5 mAb (Eculizamab) for treating paroxysmal nocturnal hemoglobinuria (PNH) found only sporadic concern of Neisseria infection. Eculizamab and PMX205 both approaches still leave intact C3-opsonization, thereby minimizing the possibility of bacterial infections. Secondly, with the C5aR1 antagonist, the downstream cascade will have the ability to form the membrane attack complex (MAC) in contrast to Eculizumab (66-68). Thus, preventing negative consequences, including Neisseria infection. While a couple of studies show an issue with the long-term blocking of C5a (67,69). It would also be important to stress that PMX205 treatment for cognitive decline post-RT would not be long-lasting and can be monitored in the clinical setting. Human clinical trials with C5aR1 inhibitors (PMX53, CCX168) for rheumatoid arthritis and ANCA-associated vasculitis showed excellent safety profiles and lack of toxicity(13,70), and Avacopan (Tavneos, CCX168) is now FDA-approved for autoimmune disease treatment. However, it has shown very limited penetration into the CNS, including the brain and spinal cord (14). To address this issue, we selected PMX205, which has shown superior brain penetration and oral availability and is neuroprotective in mouse models of neurodegenerative conditions, including AD and ALS (8,11,71). In recent clinical trials with healthy individuals and ALS patients, PMX205 did not show toxicity, drug-related adverse events, or infections (reference ACTRN12622000927729). Thus, pre-clinical mouse and human clinical trial data suggest that inhibition of the C5a-C5aR1 signaling axis is not harmful, emphasizing the translational feasibility of this approach to improve the QOL for millions of cancer survivors.

## Supporting information

All_Supplemental_Figures_Methods

## LIMITATIONS OF THE STUDY

We provide proof of the principle that C5a-C5aR1 axis inhibition via either genetic ablation of C5aR1 or pharmacologic inhibition using PMX205 reverses the detrimental neuroinflammatory and neurocognitive consequences of cranial RT. We showed this using pre-clinical cancer-free and brain cancer-bearing mouse models. Cranial RT in the clinic is often scheduled as a fractionated or hypofractionated dosing regimen in combination with temozolomide. Thus, the translational feasibility of PMX205 should also be tested using a fractionated cranial RT paradigm to approximate the biological effective dose (B.E.D.) regime for late RICD. A fractionated RT regime with concomitant and adjuvant temozolomide is expected to eliminate brain cancer and allow the study of cellular and molecular endpoints of microglial, astrocytic, and synaptic function *in vivo*. A longitudinal cognitive function assessment before and post-RT and after completion of C5aR1 inhibitor treatment would allow the restoration of cognitive function in the injured brain to be studied. Albeit, such cognitive testing may lead to fatigue in animals and limit testing of invasive tasks. Nonetheless, these proposed experiments will determine the long-term safety and neuroprotective efficacy of C5aR1 inhibitors for cranial RT-induced normal tissue toxicity.

## AUTHOR’S DISCLOSURES

The authors declare no relevant conflicts of interest or financial relationships.

## AUTHOR’S CONTRIBUTIONS

Conception and design: MMA, AJT, JEB

Development of methodology: RPK, AHD, SMK, LA, MM, ARV, MTU, SM

Acquisition of data: RPK, AHD, SMK, LA, MM, ARV, MTU, SM

Analysis and interpretation of data: RPK, AHD, SMK, JEB, MMA

Writing, review and/or revision of the manuscript: RPK, AHD, JEB, TMW, AJT, MMA

Administrative, technical, or material support: RJC, TMW, AJT, JEB, MMA.

Study supervision: RPK, JEB, MMA.

## ACKNOWLDEGMENTS

This work was supported by the National Institute of Health (NIH) awards (R01CA251110, R01CA262213, and R01CA276212), American Brain Tumor Association Discovery award (DG2000029), HESI-Thrive award (Health and Environmental Sciences Institute, 2021), and the University of California Irvine (UCI) Chao Family Comprehensive Cancer Center (CFCCC) Pilot award to M.M.A. T.M.W. is supported by the National Health and Medical Research Council (APP2009957). We also thank the UCI CFCCC Genomics Research and Technology Hub (GRTH), supported by the NIH award (P30CA062203). The content is solely the responsibility of the authors and does not necessarily represent the official views of the NIH.

## Notes

https://www.ncbi.nlm.nih.gov/geo/query/acc.cgi?acc=GSE282058

### Competing Interest Statement

The authors have declared no competing interest.

### Summary of Updates

Additional experiments and data are included to confirm our hypothesis on the neuroprotective impact of inhibiting the C5a/C5aR1 axis in reversing cognitive impairments following cranial radiotherapy for brain cancers.

