## Supplementary material for "C5aR1 inhibition alleviates cranial radiation-induced cognitive decline": All_Supplemental_Figures_Methods

**Supplemental Figure S1**

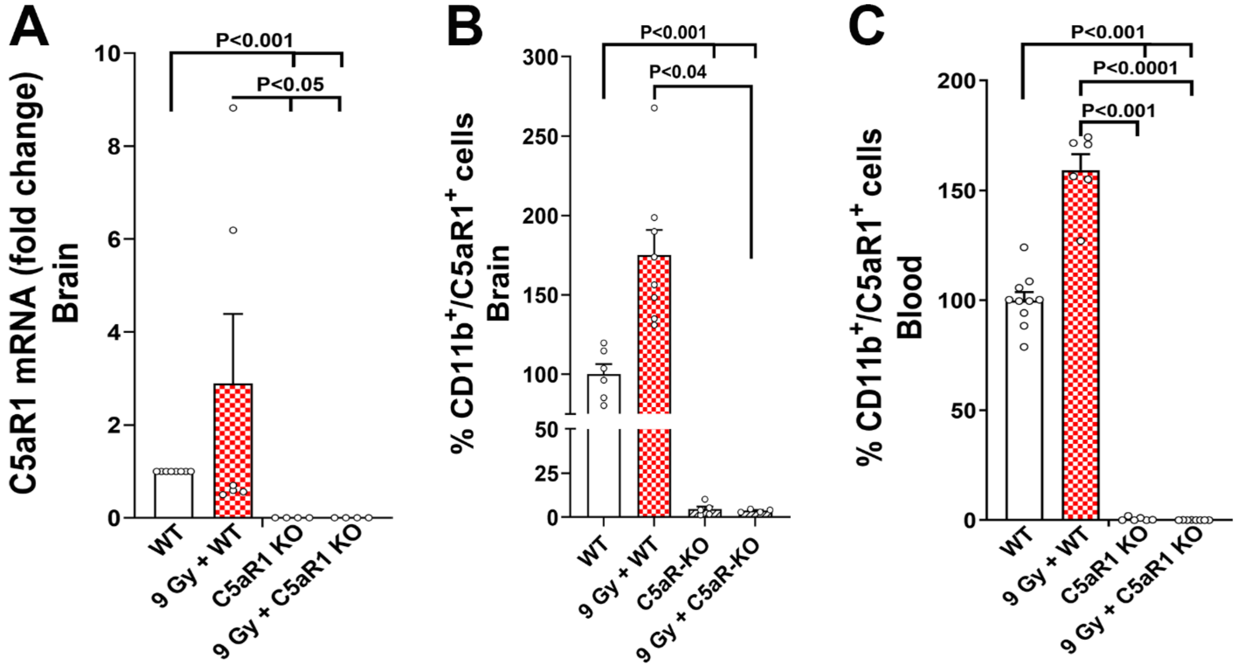


**Supplemental Figure S1:** Characterization of C5aR1 expression in the WT and C5aR1 KO mice brains (**A-B**) and blood (**B**). (**A**) qPCR for C5aR1 mRNA expression showed nearly complete ablation of C5aR1 transcript in the 0 Gy and 9 Gy irradiated brains of C5aR1 KO mice. **(B-C)** Flow cytometry analysis of C5aR1 from brains and blood also showed nearly complete ablation of C5aR1 *in vivo*. Mean ± SEM (*N*=6-10 mice per group). *P* values were derived from two-way ANOVA and Bonferroni's post hoc test.

**Supplemental Figure S2**

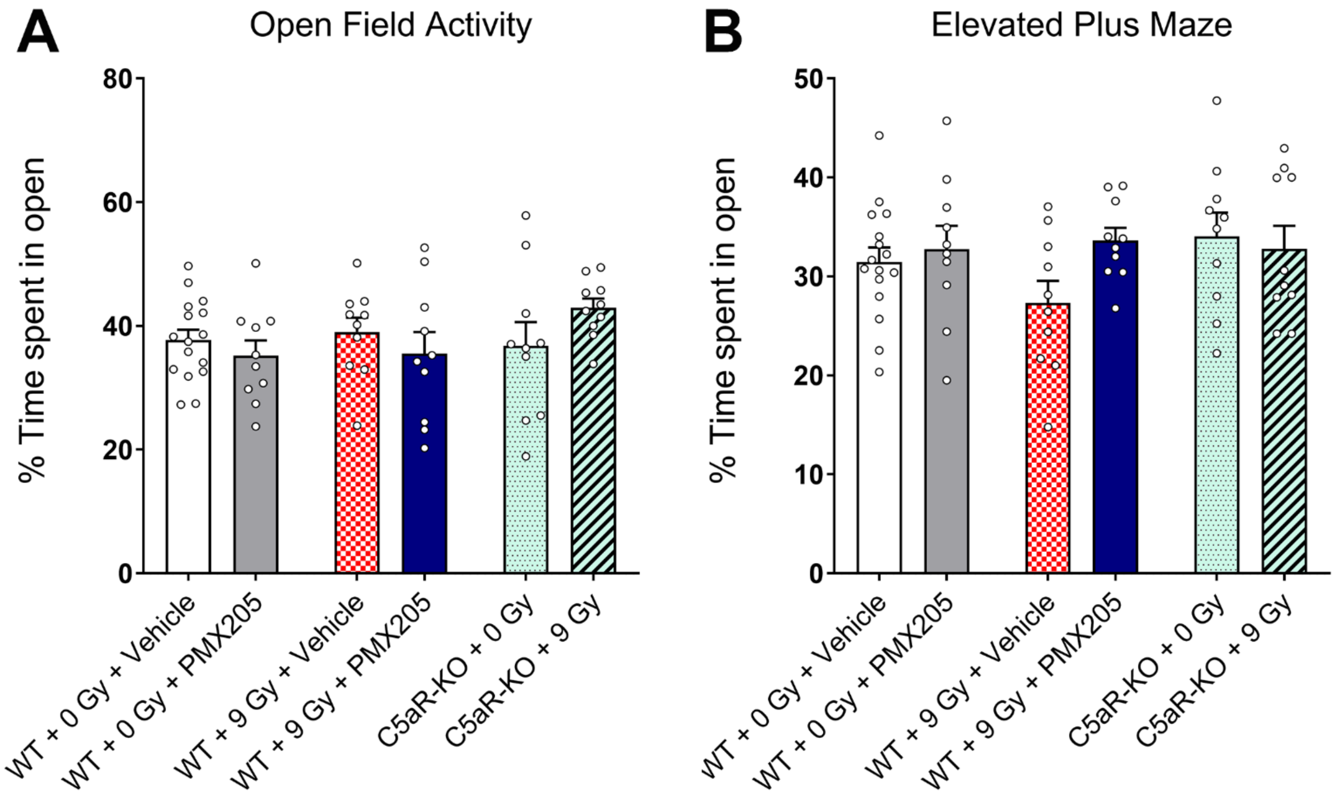


**Supplemental Figure S2.** Cranial irradiation, PMX205 treatment and C5aR1 KO did not affect the open field activity and anxiety-like behavior. No significant differences were observed between littermate 0 Gy or 9 Gy irradiated WT, WT mice treated with PMX205 and C5aR1 KO mice on the open field activity (percent time spent in open or central zone of an arena) and elevated plus maze (percent time spent in open arms) tests one-month post-irradiation. Data are presented as mean ± SEM (*N*=8-16 mice per group).

**Supplemental Figure S3**

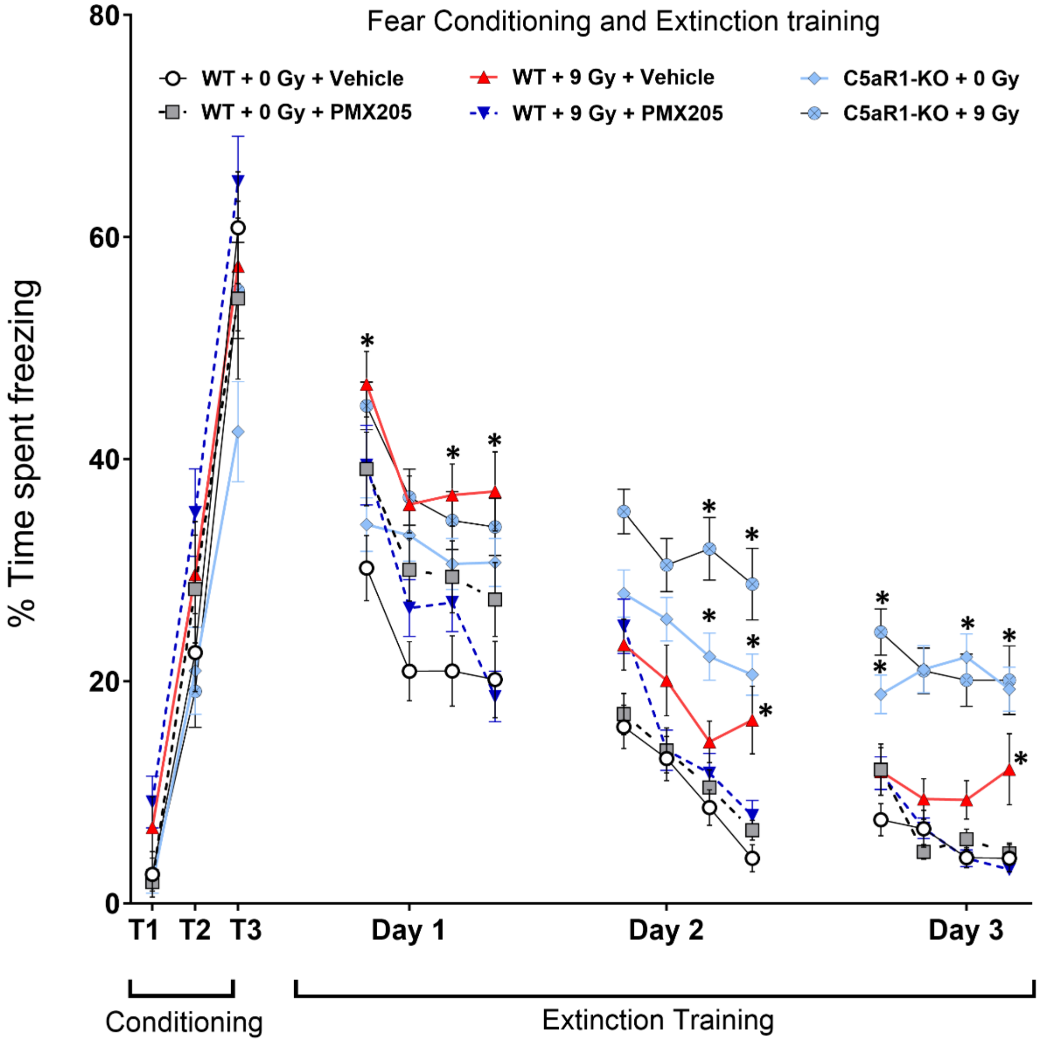


**Supplemental Figure S3.** Treatment with PMX205 or C5aR1 knockout did not impair the acquisition of conditioned fear memory, indicated by increased freezing behavior following three-tone and shock pairings (80 dB, 0.6 mA, **T1–T3**). At 24 h later, extinction training was administered every 24 h (20 tones) for 3 days. Each data point for Days 1-3 is presented as the average percentage time freezing for 5 tones (4 data points per day). All mice showed a gradual decrease in freezing behavior (**Days 1–3**). However, C5aR1-KO + 0 Gy and 9 Gy groups spent significantly more time freezing than Control + Vehicle mice. For Days 2 and 3, WT + 9 Gy + Vehicle mice also showed increased freezing compared to the Con + Vehicle. The extinction test was administered 24 h after the extinction training (three tones, no shock, as in **Fig. 1E**). WT + 9 Gy + Vehicle mice showed elevated freezing. Mean ± SEM (*N* = 10-16 mice per group).

**Supplemental Figure S4**

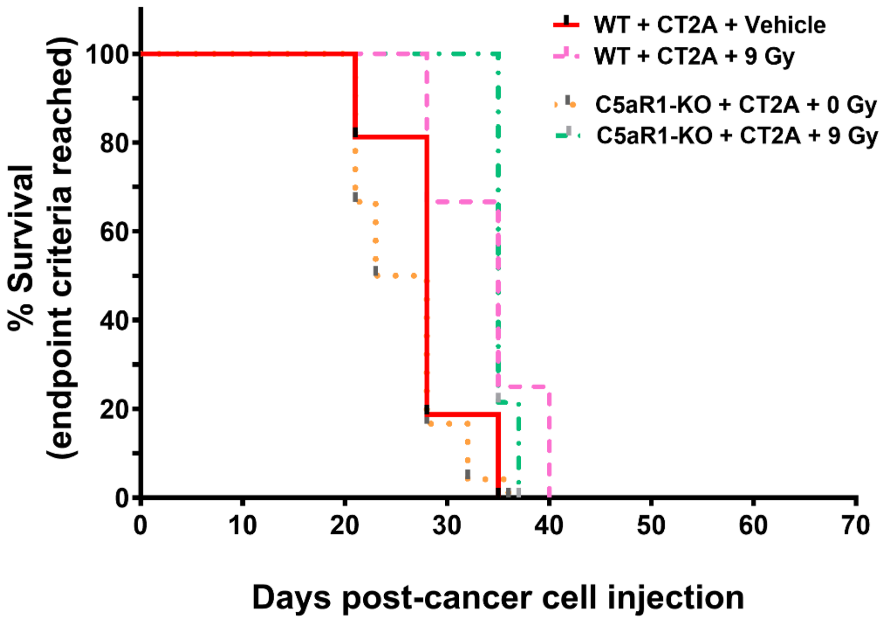


**Supplemental Figure S4.** Kaplan-Meier estimates derived from **Fig. 4E**, showing four groups (WT + CT2A ± 9 Gy, and C5aR1-KO + CT2A ± 9 Gy). CT2A$^{+}$ glioma/astrocytoma showed comparable survival estimates for all groups of mice (*N*=12-24 mice per group).

### Supplemental Figure S5

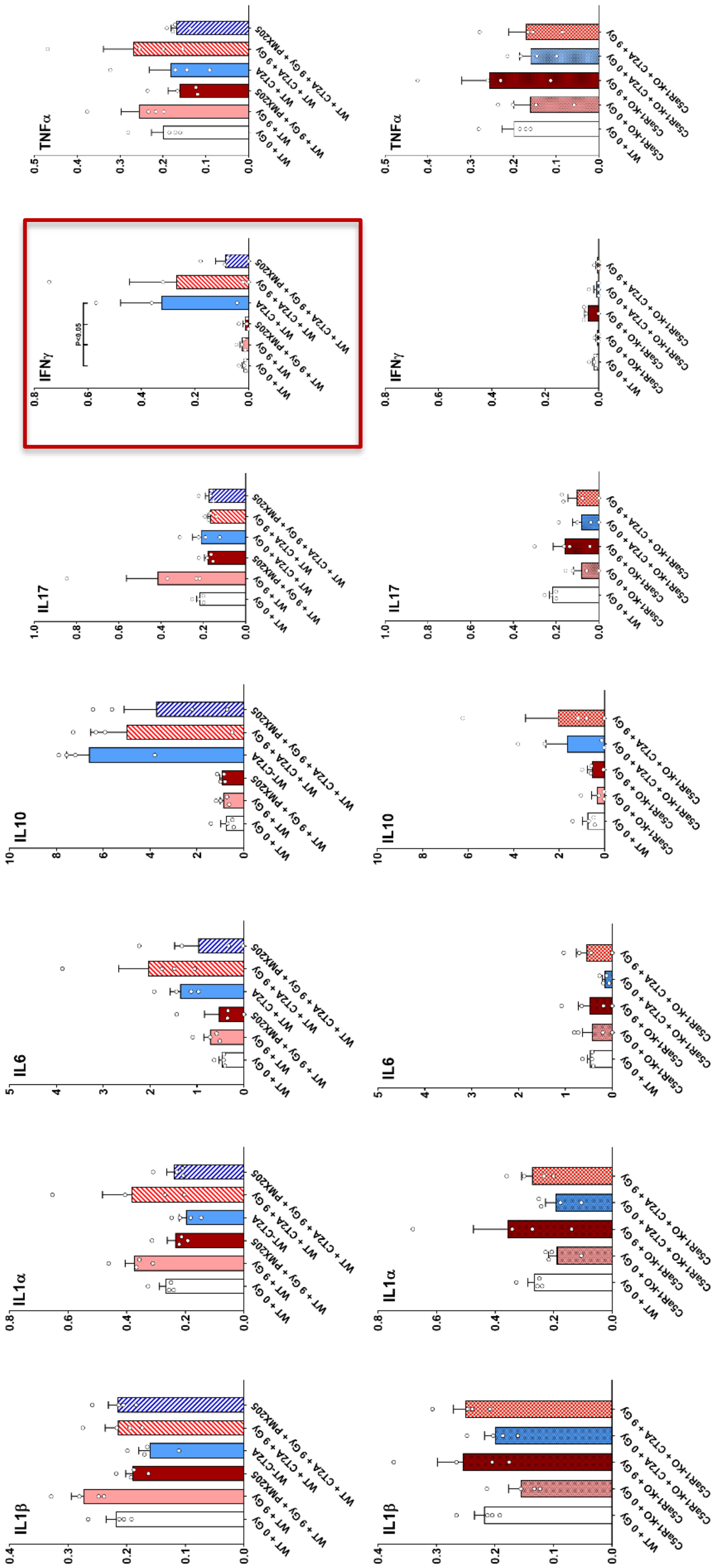


**Supplemental Figure S5.** Multiplex ELISA-based cytokine analysis of mice hippocampal extracts. No statistical significance (P≤0.05) was observed except for elevated IFNγ (red box) in the WT + CT2A and WT + CT2A + 9 Gy brain compared to WT + 0 Gy + Vehicle group. Top row, WT, and PMX205 comparisons; bottom row, WT, and C5aR1-KO comparisons. Mean ± SEM (*N* = 3-4 mice per group).

**Supplemental Figure S6**

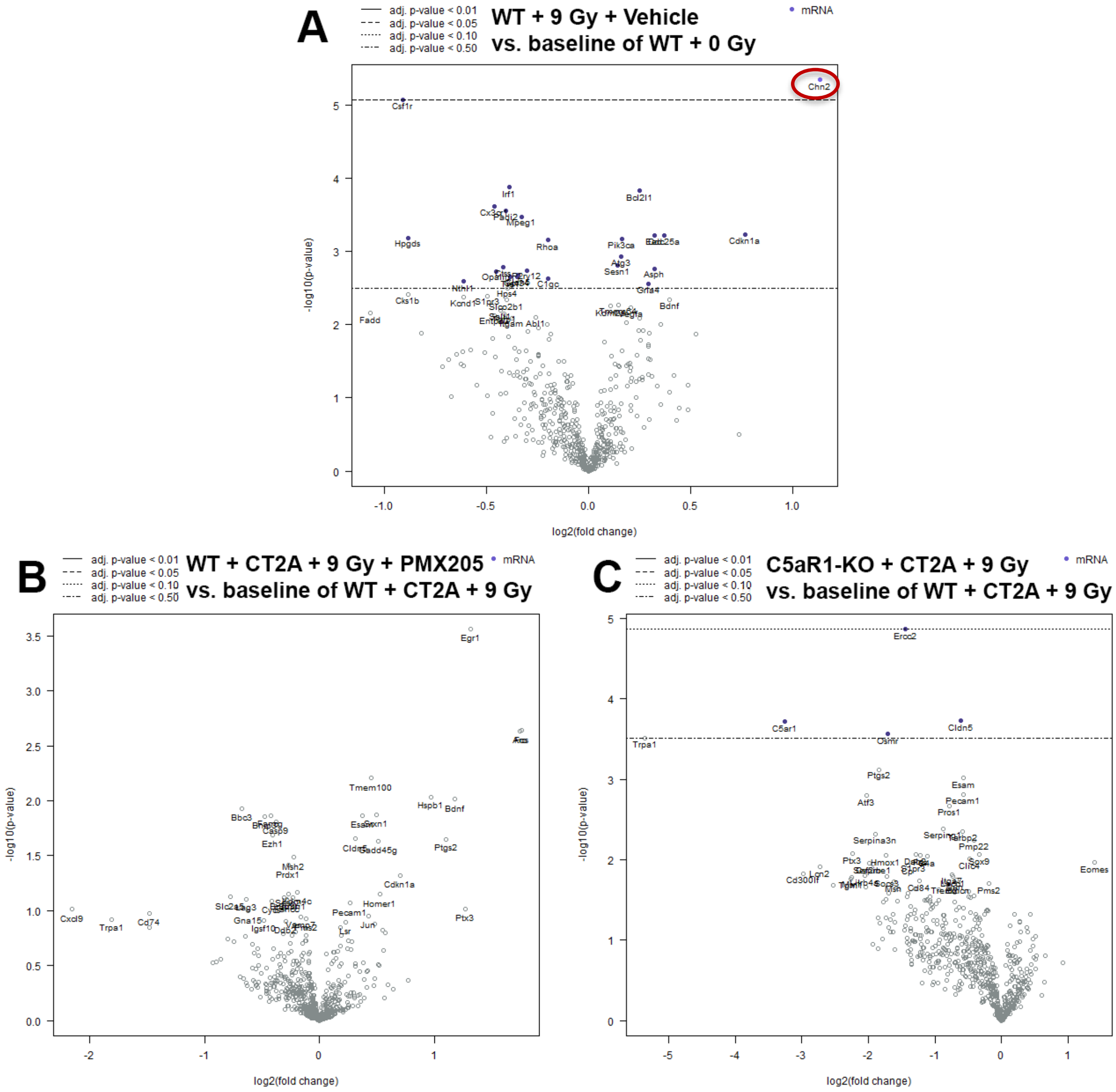


**Supplemental Figure S6.** Volcano plots of differentially expressed genes (DEGs) comparing WT + 9 Gy + Vehicle to the WT + Gy group **(A)**, WT + CT2A + 9 Gy + PMX205 **(B)**, and C5aR1-KO + CT2A + 9 Gy **(C)** to the WT + CT2A + 9 Gy group as a baseline. The log2 fold change (logFC) is plotted against the -log10 for each gene that met the minimal count threshold. No statistical significance (P≤0.05) was observed except for *Chn2* (microglial inflammation and synaptic elimination roles) in the WT + 9 Gy + Vehicle brain (P<0.05, **A**). Adjusted P values (P≤0.05) were derived from the Benjamini-Yekutieli False Discovery Rate (FDR) method (N = 4 brains/group).

### Supplemental Table T1

#### WT+ 9 Gy + PMX205 *vs.* WT + 9 Gy + Vehicle

| mRNA | Affilated Gene Sets |
|---|---|
| Apc | Apoptosis, Astrocyte Function, Wnt |
| Prkce | Growth Factor Signaling, Autophagy |
| Pten | Growth Factor Signaling, Autophagy, Adaptive Immune Response, DNA Damage, Lipid Metabolism |
| Ikbkb | Growth Factor Signaling, Innate Immune Response, Adaptive Immune Response, Apoptosis, Cytokine Signaling, Inflammatory Signaling, NF-kB |
| Grin2a | Growth Factor Signaling, Neurons and Neurotransmission, Adaptive Immune Response, Cytokine Signaling, Insulin Signaling |
| Grin2b | Growth Factor Signaling, Neurons and Neurotransmission, Adaptive Immune Response, Cytokine Signaling, Insulin Signaling |
| Cx3cl1 | Innate Immune Response, Cytokine Signaling, Inflammatory Signaling |
| Cldn5 | Matrix Remodeling |
| Abcc3 | Microglial Function |
| Gria2 | Neurons and Neurotransmission |
| Homer1 | Neurons and Neurotransmission |
| Dlg1 | Neurons and Neurotransmission, Autophagy |
| Calr | Neurons and Neurotransmission, Autophagy, Adaptive Immune Response |
| Bag3 | Apoptosis, Cellular Stress |
| Agt | Astrocyte Function |
| Lamp2 | Autophagy |
| Pink1 | Autophagy, Cellular Stress |
| Gstm1 | Cellular Stress, Inflammatory Signaling |
| Bcl2l1 | Growth Factor Signaling, Autophagy, Innate Immune Response, Apoptosis, Cytokine Signaling, NF-kB |
| S100b | Growth Factor Signaling, Innate Immune Response, Inflammatory Signaling, Astrocyte Function |
| Nfkbia | Growth Factor Signaling, Innate Immune Response, Adaptive Immune Response, Apoptosis, Inflammatory Signaling, NF-kB |
| Prkaca | Growth Factor Signaling, Neurons and Neurotransmission, Autophagy, Adaptive Immune Response, Apoptosis, Cytokine Signaling, Wnt, Lipid Metabolism |
| Mag | Growth Factor Signaling, Oligodendrocyte Function, Matrix Remodeling |
| Asph | Microglial Function |
| Bola2 | Microglial Function |
| Cd83 | Microglial Function |
| Cox5b | Microglial Function |
| F3 | Microglial Function |
| Pmp22 | Microglial Function |
| Rpl28 | Microglial Function |
| Rps3 | Microglial Function |
| Slc2a1 | Microglial Function |
| Tmcc3 | Microglial Function |
| Lamp1 | Microglial Function, Autophagy |
| Spp1 | Microglial Function, Matrix Remodeling |
| Csf1 | Microglial Function, Growth Factor Signaling, Autophagy, Innate Immune Response, Cytokine Signaling, Astrocyte Function |
| Lfng | Microglial Function, Notch |
| Plekhb1 | Neurons and Neurotransmission |
| Slc17a6 | Neurons and Neurotransmission |
| Gja1 | Neurons and Neurotransmission, Astrocyte Function |
| Bcas1 | Oligodendrocyte Function |
| Cnp | Oligodendrocyte Function |
| Mobp | Oligodendrocyte Function |
| Mog | Oligodendrocyte Function |
| Myrf | Oligodendrocyte Function |
| Plp1 | Oligodendrocyte Function |

Upregulated    Downregulated

#### C5aRKO + 9Gy *vs.* WT + 9 Gy + Vehicle

| mRNA | Affiliated Gene Sets |
|---|---|
| Hprt | Apoptosis |
| Hrk | Apoptosis |
| Mfge8 | Autophagy |
| Atg9a | Autophagy, Cellular Stress |
| Map1lc3a | Autophagy, Cellular Stress |
| H2afx | Cellular Stress, Epigenetic Regulation, Wnt, Cell Cycle |
| Kdm5b | Epigenetic Regulation |
| Mbd3 | Epigenetic Regulation |
| Smarcd1 | Epigenetic Regulation |
| Prkar2b | Growth Factor Signaling, Apoptosis, Cell Cycle |
| Rhoa | Growth Factor Signaling, Autophagy, Angiogenesis, Wnt |
| Bad | Growth Factor Signaling, Autophagy, Cellular Stress, Adaptive Immune Response, Apoptosis |
| Sqstm1 | Growth Factor Signaling, Autophagy, Cytokine Signaling |
| Dlg4 | Growth Factor Signaling, Neurons and Neurotransmission, Angiogenesis, Adaptive Immune Response, Signaling, Insulin Signaling |
| Prkacb | Growth Factor Signaling, Neurons and Neurotransmission, Autophagy, Angiogenesis, Adaptive Immune Response, Apoptosis, Wnt, Lipid Metabolism |
| Akt2 | Growth Factor Signaling, Neurons and Neurotransmission, Autophagy, Innate Immune Response, Apoptosis, Wnt, Insulin Signaling, Carbohydrate Metabolism |
| Rac1 | Growth Factor Signaling, Neurons and Neurotransmission, Autophagy, Innate Immune Response, Adaptive Immune Response, DNA Damage, Wnt |
| Lingo1 | Growth Factor Signaling, Oligodendrocyte Function |
| C1qa | Innate Immune Response |
| C1qb | Innate Immune Response |
| C1qc | Innate Immune Response |
| Cx3cl1 | Innate Immune Response, Cytokine Signaling, Inflammatory Signaling |
| Olfml3 | Matrix Remodeling |
| Ak1 | Microglia Function |
| Clstn1 | Microglia Function |
| Fscn1 | Microglia Function |
| Ldha | Microglia Function |
| Man2b1 | Microglia Function |
| P2ry12 | Microglia Function |
| Pacsin1 | Microglia Function |
| Ptms | Microglia Function |
| Rps2 | Microglia Function |
| Rps21 | Microglia Function |
| Trem2 | Microglia Function, Adaptive Immune Response, Inflammatory Signaling |
| Cx3cr1 | Microglia Function, Cytokine Signaling |
| Stmn1 | Microglia Function, Growth Factor Signaling |
| Map2k1 | Microglia Function, Growth Factor Signaling, Autophagy, Innate Immune Response, Angiogenesis, Adaptive Immune Response, Cytokine Signaling, Wnt, Insulin Signaling |
| Kcnd1 | Microglia Function, Neurons and Neurotransmission |
| Apoe | Microglia Function, Neurons and Neurotransmission, Cellular Stress, Astrocyte Function, Lipid Metabolism |
| Islr2 | Neurons and Neurotransmission |
| Nrgn | Neurons and Neurotransmission |
| Rbfox3 | Neurons and Neurotransmission |
| Slc17a7 | Neurons and Neurotransmission |
| Syp | Neurons and Neurotransmission |
| Grm3 | Neurons and Neurotransmission, Astrocyte Function |
| Nlgn2 | Neurons and Neurotransmission, Matrix Remodeling |
| Eed | Cellular Stress, Epigenetic Regulation |
| Ercc2 | DNA Damage, Inflammatory Signaling |
| Kat2b | Epigenetic Regulation, Notch |
| Dst | Microglia Function, Matrix Remodeling |
| Gria4 | Neurons and Neurotransmission |

### SUPPLEMENTAL METHODS AND MATERIALS

#### Resource Availability

#### Materials availability

This study did not generate any new unique reagents.

#### Data and code availability

- All data reported will be shared by the lead contact upon a reasonable request.
- This article did not report the original code.
- Any additional information required to reanalyze the data reported in this article is available from the lead contact upon a reasonable request.
- The NanoString Neuroinflammation gene expression panel data is available via the NIH Gene Expression Omnibus (GEO) archive, reference number: GSE282058.

### EXPERIMENTAL MODEL AND SUBJECT DETAILS

#### Mice

All animals used in this study were cared for in accordance with NIH guidelines and approved by the Institutional Animal Care and Use Committee (IACUC) at the University of California, Irvine.

Wild-type male mice, aged 15-16 weeks (C57BL/6J, RRID:IMSR_JAX:000664) were obtained from The Jackson Laboratory and housed in standard conditions (20 °C±1 °C; 70% ± 10% humidity;12h:12h light and dark cycle) in groups of 2-5 mice per cage. Homozygous C5aR1 KO males (C5aR -/-), aged 15-16 weeks, were bred in-house. The transgenic mice pair were graciously provided by Dr. Rick A. Wetsel, University of Texas Health Science Center, Houston. Dr. Wetsel had previously developed a C5aR -/- transgenic mouse model via targeted deletion of the C5a receptor 1 gene (PMID: 18063050). The in-house breeding colony was maintained and expanded via selective inbreeding to produce homozygous offspring, and the genotype of digested tail clips or ear punches was confirmed by PCR. All C5aR1 KO mice were housed in standard conditions (20 °C±1 °C; 70% ± 10% humidity;12h:12h light and dark cycle) in groups of 2-5 mice per cage. For both wild-type and C5aR1 KO, littermates were randomly assigned to experimental groups. Groups that underwent intracranial tumor induction and/or cranial radiation therapy (RT).

### METHOD DETAILS

#### Experimental design

C5a receptor inhibition and its effects on cognition in the irradiated brain were first studied in non-cancer-bearing mouse models, either with pharmacological intervention via cyclic hexapeptide C5a inhibitor PMX205 or through genetic ablation with the C5aR1 Knockout (KO) transgenic mouse model. For this study, 15-16 weeks male mice were divided into the following groups: 1) WT mice receiving 0 Gy or 9 Gy cranial radiation therapy (CRT) with or without PMX205 treatment and 2) C5aR1 KO mice receiving 0 Gy or 9 Gy CRT. One month post-treatment, cognitive function testing was performed, followed by euthanasia and tissue harvesting. For the tumor-bearing model, WT C57Bl6 male mice (15-16 weeks old) were divided into the following groups and applied to both the astrocytoma (CT2A-Luc) and glioblastoma (GL261) tumor types: Tumor + Vehicle, Tumor + RT, Tumor + PMX205, and Tumor + RT + PMX205. Tumor growth and animal health were monitored over the period of one- to two-month post-treatment. A cohort of mice underwent cognitive function testing at three- to four weeks post-treatment.

#### Preparation of mouse glioma cells for tumor induction

CT2A-Luc mouse astrocytoma cells (Sigma-Aldrich, Cat. SCC195, RRID:CVCL_ZJ60 ) and GL261 mouse glioblastoma cells (NCI-DTP, Cat. Glioma 261, RRID:CVCL_Y003) were cultured in monolayer with Gibco™ DMEM (1X) supplemented with GlutaMAX™-I (FisherSci, Cat. 10-569-010) and 10% Gibco™ FBS (FisherSci, Cat. 10-082-147) at 37°C and 5% $CO_2$. For surgical preparation, CT2A-Luc and GL261 cell cultures were first trypsinized with Gibco™ TrypLE™ Express Enzyme (FisherSci, Cat. 12-605-010), spun down at 300g for 5 minutes, resuspended in Gibco™ Hibernate™-A Medium (FisherSci, Cat. A1247501) and kept on ice until the implantation. Successful tumor induction in mouse subjects required a minimal cell 2,000 cells per brain, injected in a total volume of 2 µl per animal.

#### Stereotaxic intracranial Implantation of mouse glioma cells

Experiment groups undergoing tumor induction were first anesthetized with 2.0% v/v inhalant isoflurane (Dechra) delivered at 1000 cc/min ± 10% with a passive scavenging anesthesia system (VetEquip, Cat. 922100). Mice were subsequently shaved with an electric trimmer, mounted to a digital stereotaxic rig (Stoelting, Cat. 51730UD) and kept under anesthesia with 2.0% inhalant isoflurane at 500 cc/min ± 10%. The shaved scalp was prepped with Betadine® Surgical Scrub (Patterson Veterinary, Cat. 07-836-3379) followed by Sterile Alcohol Prep Pad (FisherSci, Cat. 06-669-62); this was repeated two more times. A single midline sagittal incision was made with a scalpel (FisherSci, Cat. 08-927-5B) and the underlying soft tissue was treated with 3% hydrogen peroxide (FisherSci, Cat. H325-500) to expose bregma, which is the origin point for stereotaxic coordinates in mice. The coordinates for the tumor cell implantation were anterior-posterior (AP) +1.54 mm, medial-lateral (ML) +2.00 mm, and dorsal-ventral (DV) -3.00 mm. The AP-ML coordinates were demarcated with a fine-tip pen, and a bone drill was used to create an opening in the cranium. A 10 µl Hamilton syringe with a cemented needle (Hamilton, Cat. 80300) was used to draw up 2 µl of the tumor cell suspension. The dose was injected at DV -3.00 mm in two stages, with a 60-second pause after the first microliter and a six-minute pause after the second microliter. The needle was then withdrawn in a stepwise manner from DV -2.00 mm to -1.00 mm, with a 60-second pause at each step to prevent the backflow of the tumor suspension. The incision was closed in an interrupted pattern with size 6-0 polyglactin-coated sutures (Patterson Veterinary, Cat. 07-891-0161) and tissue adhesive (Patterson Veterinary, Cat. 07-805-5031). Animals were placed in a recovery cage on a heating pad and monitored until they gained consciousness, during which animals received a 1.0 ml intraperitoneal injection of saline (0.9% NaCl, Patterson Veterinary, Cat. 07-800-9424). Post-surgical analgesic care consisted of a subcutaneous injection of Buprenorphine HCl (MWI Health) diluted in saline to 0.1 mg/kg body weight. This was delivered once at the end of the surgery and then again at 12 hours later.

**Tumor growth measurements for the CT2A-Luc- and GL261-implanted animals**

Astrocytoma/glioma (CT2A-Luc) tumor growth was measured via bioluminescence imaging (BLI). Animals received a 200 µl intraperitoneal injection of the bioluminescent reporter substrate luciferin, which was prepared as a 15 mg/ml stock solution from D-Luciferin, sodium salt (GoldBio, Cat. eLUCNA) and Gibco™ DPBS (no calcium, no magnesium, FisherSci, Cat. 14-190-144). Injected mice were then anesthetized using isoflurane (as above) and placed in a prone position inside the IVIS Lumina III *in vivo* Imaging System (Revvity, Perkin Elmer, Cat. CLS136334). The enzymatic activity of the luciferase expressed by the CT2A-Luc cells was captured, visualized, and quantified with the accompanying Living Image Software (Revity, Perkin Elmer, Cat.128110). The time from injection to image capture was optimized by performing serialized acquisitions on two subjects until a peak emittance was observed at approximately 12-14 minutes post-injection. This time interval was maintained for all subsequent subjects and weekly time points.

Glioblastoma tumor growth was tracked using cone-beam computed tomography (CBCT) using the SmART+ image-guided X-ray research irradiator (Precision, Inc.). To enhance soft-tissue contrast, 50 mg of iohexol (Omnipaque™ 350 mgI/ml, GE Healthcare) was injected retroorbitally into anesthetized mice immediately prior to imaging. DICOM images were analyzed with OsiriX Lite Software (Pixmeo SARL) to calculate the tumor size.

**Cranial radiation therapy (RT)**

RT was administered to the WT and C5aR1 KO mice, to the assigned tumor groups approximately 7-9 days post-tumor induction using the SmART+ X-ray irradiator. A total of 9 Gy was delivered using a 0.3mm Copper filter, a 10×10 mm fixed collimator, and a photon spectrum of 225 kVp at 20 mA for 1 minute and 50 seconds. The treatment was delivered in 2 equal doses of 4.5 Gy in series on the same day in the sagittal plane to minimize damage to tissue outside the region of interest.

**PMX205 preparation and administration**

The cyclic hexapeptide C5a inhibitor PMX205 was manufactured at Mimotopes (Australia) in consultation with Drs. Trent Woodruff and Richard Clark, The University of Queensland, Brisbane. The drug IC50 activity was confirmed in the Woodruff lab before shipping to the U.S. An initial 10 mg/ml stock solution comprised 50% DMSO (FisherSci, Cat. AAJ66650K2) in Milli-Q $H_2O$. The PMX205 treatment paradigm consisted of one week of daily subcutaneous injections at 1 mg/kg body weight and a month-long consumption of 100 µg/day (or 20 µg/ml) in the drinking water, both of which began 48 hours post-CRT. Water consumption was tracked weekly, and bottles were rotated thrice weekly to encourage complete fluid intake.

**Object recognition memory (ORM) and object location memory (OLM)**

Object recognition memory (ORM) and object location memory (OLM) tests depend on intact hippocampal, perirhinal cortex, and medial prefrontal cortex functions (2,3). The ORM task evaluates episodic memory by measuring animal's preference to explore novel objects, whereas the OLM task evaluates spatial recognition. For both tasks, the setup includes testing rooms equipped with appropriate lighting (~50 Lux), four square arena boxes, camera recording hardware (Noldus), and tracking software (EthoVision XT 17). The ORM and OLM tasks were conducted as described previously (1 and 4, respectively). Briefly, for the ORM task, the mice were habituated in open field arena boxes (33 × 33 × 33 cm) with a thin layer of bedding and without any objects for 10 minutes each run in 3 consecutive days. For the ORM task, the familiarization phase allowed the mice to explore two identical plastic toys that were magnetically secured in place 16 cm apart from opposing corners for 5 minutes. After a 5-minute interval in their cages, a novel object replaced one of the familiar objects (both toys were thoroughly cleansed with 10% ethanol and dried). Mice were then returned to the arena boxes and allowed to explore for 5 minutes. Following the test phase, all mice were returned to their home cage. The OLM task was conducted the following week after the ORM task. Mice were allowed to familiarize themselves with the testing room and open field arena boxes (33 × 33 × 33 cm) with a water-resistant, matte finish blue tape down the inside wall of the arena boxs and a layer of bedding. Mice were allowed to explore two identical toys that were placed 16 cm apart from each other for 5 minutes. Following similar steps from the ORM task, mice were returned to the home cage for 5 minutes while the objects were cleansed with 10% ethanol. One object was then moved to a new location inside the box (novel place), 16 cm from the opposite corner of the other toy, which remained in its former spatial location (familiar phase). Mice were returned to arena boxes and allowed to explore for 5 minutes. Mice behavior from all phases was recorded, and the "head direction to zone" function from Ethovision software was utilized to track the exploration time. Furthermore, to ensure unbiased analysis, time spent interacting (nose within 2 cm) with familiar vs novel (or relocated) objects was scored by observers blinded to the experimental conditions. The discrimination index (DI) was calculated for each animal using the equation: ([Novel object or location exploration time/Total exploration time] – [Familiar object or location exploration time/Total exploration time]) × 100.

**Elevated plus maze (EPM)**

The EPM test is a measure of anxiety that consists of two bisecting platforms in the shape of a plus sign constructed some distance off the ground, with one open arm with no walls or roof and one closed arm that consists of high barriers and a darker environment (7). Prior to testing, the floor and walls of the maze were cleaned with odor-elimination disinfectant (Virkon, University Laboratory Animal Resources, UCI) and then dried with paper towels. Each mouse subject was placed at the intersection of both arms and allowed to explore for a duration of 5 minutes. Testing was recorded with the same camera hardware and analyzed with accompanying tracking software (Noldus). The ratio of time spent in the open versus closed arms was then compared across experimental groups as a proxy for anxious behavior.

**Fear extinction memory**

To determine if C5a inhibition or knockout in the irradiated brain affects amygdala-hippocampal circuit-dependent fear conditioning learning and fear memory consolidation, we performed fear extinction behavior reliant on hippocampal function (5,6). Testing occurred in a behavioral conditioning chamber (17.5 × 17.5 × 18 cm, Coulbourn Instruments) with parallel steel grid shock floors (3.2 mm diameter slats, 8 mm spacing), and a waste collection tray sprayed with 10% vinegar. For the initial fear conditioning phase (day 1), mice were allowed to habituate to the chamber for two minutes. Three pairings of an auditory conditioned stimulus (16 kHz tone, 80 dB, lasting 120 sec; CS) co-terminating with a foot shock unconditioned stimulus (0.6 mA, 1 sec; US) were presented at two-minute intervals. On the following three days of extinction training (days 2-4), mice were initially habituated to the same context for two minutes before being presented with 20 non-US reinforced CS tones (16 kHz, 80 dB, lasting 120 sec, at 5 sec intervals). On the final day of fear testing (day 5), mice were presented with only three non-US reinforced CS tones (16 kHz, 80 dB, lasting 120 sec) at two-minute intervals in the same context. Freezing behavior was recorded with a camera mounted above the chamber and scored by an automated measurement program (FreezeFrame, Coulbourn Instruments). FreezeFrame algorithms calculated a motion index for each frame of the video, with higher values representing greater motion. An investigator blinded to the experimental groups set the motion index threshold representing immobility for each animal individually based on identifying a trough separating low values during immobility and higher values associated with motion. Freezing behavior was defined as continuous bouts of one second or more of immobility. The data for the percentage of time each mouse spent freezing was then calculated for the final day of the extinction test.

**Isolation of immune cells from brains and blood**

On the day of analysis, mice blood and whole brains were freshly collected. Mice were deeply anesthetized with 2% isoflurane v/v (inhalation, Dechra) prior to the terminal blood collection and intracardiac perfusion. The heart was exposed after the opening of the thoracic cavity, and blood was carefully drawn via cardiac puncture (volume ~ 200uL) using a 28G needle and immediately placed in tubes coated with saline (0.9% NaCl, Sigma-Aldrich Cat. S9888) containing 20 U/ml heparin (Sigma-Aldrich, Cat. H3149) in ultra-pure water (Milli-Q $H_2O$) at room temperature (rt) for later processing. Next, intracardiac perfusion using a peristaltic pump was performed using ice-cold saline with 10 U/ml of heparin (Sigma-Aldrich) to prevent blood clotting. Brains and spleens were immediately collected and placed in separate tubes filled with Gibco$^{TM}$ HBSS (FisherSci, Cat. 14-175-079) with 10% Gibco$^{TM}$ FBS (FisherSci, Cat. 10-082-147) on ice. Once all the samples were collected, brains were transferred to tubes containing digestion buffer (0.05% Collagenase D Sigma-Aldrich Cat. C0130-500MG; 0.1µM TLCK, Sigma-Aldrich Cat. T7254-100MG, 10µg/mL DNAse 1, Zymo Cat. E1011-A; 10mM HEPES, FisherSci Cat. 15-630-080 in HBSS, FisherSci) and mechanically dissociated by mincing and incubated at 37$^{o}$C for 20 minutes. Minced brain tissues were sequentially forced through a 70 µm cell strainer and centrifuged to discard the supernatant layer. Cells were then resuspended and mixed in 37% Percoll gradient (prepared Isotonic percoll by mixing Percoll, FisherSci Cat. 45001748, with 5% 10X DPBS, FisherSci Cat.14-200-075; diluted to 37% Percoll using RPMI, FisherSci Cat. 61870036, with 5% FBS, FisherSci) and centrifuged at 400g, 4ºC for 25 minutes with brake and acceleration off only for this step. Myelin from the top opaque white layer was removed and discarded. Immune cells of interest residing in the pellet at the bottom layer were washed in 1x PBS (100 mM, pH 7.4) at 300g, 4°C for 3 minutes, and the supernatants were discarded. The pellets containing cells of interest were resuspended in FACS buffer (1X DPBS, Genesee Scientific Cat. 25-508; 2% FBS, FisherSci) that were ready to be stained for flow cytometry. Spleens were crushed between two glass slides (FisherSci Cat. 22-037-246) and transferred to tubes filled with FACS buffer. Samples were centrifuged (10000g, 4ºC for 5 minutes), supernatant was discarded, and samples were incubated with red blood cell lysis buffer (FisherSci Cat. 501128916) at rt for 5 minutes. Samples were then washed, filtered through 70 µm cell strainer, centrifuged and collected from the pellet. Blood samples were incubated in the red blood cells lysis buffer in the dark for 15 minutes; samples were washed and centrifuged (500g, at rt for 5 minutes, twice) and resuspended in FACS buffer and ready to be stained for flow cytometry.

**Flow Cytometry**

Single-cell suspension from brains, blood, and spleen were divided into individual FACS tubes; samples were washed and resuspended in 30 µL of FACS buffer and incubated with antibodies. Brain and spleen cells were stained with 1 µL from each FITC anti-mouse CD45 (Biolegend, Cat. 157214), PE anti-mouse CD11b (Biolegend, Cat. 101208), APC anti-mouse CD88 (Biolegend, Cat. 135808) antibodies and DAPI (1µmol/L, FisherSci Cat. D1306) in the dark at rt for 30 minutes. Cells from blood samples were stained with 1µL from each FITC anti-mouse CD11b (Biolegend, Cat. 101206), PE anti-mouse Ly6-C (Biolegend, Cat. 128008), and APC anti-mouse CD88 (Biolegend, Cat. 135808) in the dark at rt for 20 minutes. Each sample was washed with 2mL of 1X DPBS (300g, 4ºC for 3 minutes) and then 300µL of FACS buffer was added to each tube. Samples were acquired on LSRFortessa (BD Bioscience) or a FACSCelesta (BD BioScience) flow cytometer and data was analyzed using FlowJo software (Tree Star).

**Multiplex ELISA for cytokines**

For the analysis of brain cytokines, freshly dissected hippocampus (3-4 mice/group) were homogenized with protease inhibitor cocktail made in PBS (Thermo Fisher Scientic), centrifuged (500g, at 4°C for 10 min), washed using sterile PBS (15000g, 4°C for 5 min) and supernatants were assayed for cytokines (TNFα, IL-1β, IL-6, IL-1α, IL-10, IL-17, and $IFN_{\gamma}$) using a magnetic bead-based customized multiplex ELISA (Thermo Fisher Scientific).

**RNA Extraction**

Fresh frozen brains from each group were homogenized in QIAzol Lysis Reagent (Qiagen, Cat. 79306). Total RNA were extracted and purified following precisely the recommendations in the RNeasy Plus Universal Mini Kit (Qiagen, Cat. 73404). RNA quality and quantity was evaluated using a Nanodrop (Thermo Fisher, Cat. 13-400-525), with all samples showing a wavelength 260/280 absorbance ratio between 2.0–2.1 and 260/230 ratio between 2.0–2.2. RNA integrity was further assessed using Bioanalyzer 2100 (Agilent Technologies, Waltham, MA, USA), with all samples displaying RIN > 8.5. RNA samples were stored at -80°C for further molecular assays.

**PCR Analysis**

Approximately 0.5µg of RNA from each sample were combined with master-mix (200 u/µL M-MLV Reverse Transcriptase, Promega Cat. M170A, 1.25µM RT 5X buffer, Promega Cat. M531A, 0.5mg/mL Oligo (dT) 18 Primer, FisherSci Cat. S0131, 100mM dNTP, FisherSci Cat. 10297-018, 40 u/µL RNAsin, Promega Cat. N2511, 2.5µM DTT FisherSci Cat. 707265ML in DEPC Water, FisherSci Cat. AM9920). RNA samples were reversed-transcribed to cDNA under these thermocycler conditions (Bio-rad, Cat. 1851148): 38°C for 60 minutes and 90 °C for 5 minutes. RT-PCR reactions was performed using pre-designed Taqman probes (FisherSci Cat.4453320; Assay ID: GAPDH: Mm99999915_g1; CD88:Mm00500292_s1) and Taqman Universal Master Mix, no AmpErase UNG (Thermo Fisher Scientific, Cat. 4324018), following the thermocycler parameters precisely according to the manufacturer's recommendations. Gene expression data are presented as $2^{\wedge-(\Delta\Delta Ct)}$ values.

**NanoString Neuroinflammation gene panel analysis**

We used the commercially available nCounter Mouse Neuroinflammation Panel (Nanostring, Cat. Cat. XT-CSO-MNROI1-12) to evaluate neuroinflammatory gene expression across groups. This panel profiles 757 genes, along with 13 housekeeping genes, and focuses on three main areas of neuroinflammation: immunity and inflammation, neurobiology and neuropathology, and metabolism and stress.

For each sample, 20 ng of RNA was extracted from fresh-frozen brain tissue. Gene expression was normalized using the 13 housekeeping genes included in the panel: GUSB, TBP, SUPT7L, LARS, TADA2B, CSNK2A2, CCDC127, MTO1, ASB10, XPNPEP1, FAM104A, AARS, and CNOT10, to minimize sample-to-sample variability. All procedures involving the nCounter XT Codeset Gene Expression Panel followed NanoString guidelines, with nSolver software version 4.0 used for data normalization. To ensure experiment quality, nSolver Advanced Analysis software assessed hybridization efficiency using six internal positive controls and eight negative RNA control transcripts provided in the CodeSet. These six positive controls were also utilized to normalize count variability across different runs.

**Fixed brains collection**

Mice inhaled isoflurane and were euthanized using intracardiac perfusion technique using ice-cold saline (0.9% NaCl with 10 U/ml heparin) until the liver turned slightly pale and the venous outflow was clear. Saline with heparin (10 U/ml) was then replaced with 4% paraformaldehyde (Sigma-Aldrich Cat. 158127, pH 7.4) and steadily perfused until the animal body became stiff. Whole brains were extracted and soaked in 4% PFA overnight and switched to store in PBS-0.05% sodium azide (Sigma-Aldrich Cat. S2002, pH 7.4). Brains were immersed in a sucrose gradient (10% to 30% w/v, Sigma Cat. S7903, pH 7.4) to dehydrate the brains and avoid crystal formation that could disrupt the brain structure during cryo-sectioning. Whole brains were then embedded in OCT compound (VWR Cat. 25608903) and cryo-sectioned (floating 30µm, coronal) using a cryostat (Microm Cryostar HM525 NX, Epredia, MI). Brain sections were stored in 24-well plates in PBS (100 mH, pH 7.4 with 0.05% Sodium Azide (Sigma-Aldrich, pH 7.4) for the IHC experiments.

**Immunohistochemistry**

Floating frozen brain sections were collected from each group (two sections with visible mid hippocampus region per brain, four brains per group) for immunohistochemistry. For synaptophysin, PSD95, C5aR1 and CD88-IBA1 staining, tissue sections were washed in 1x TBS (100 mM, pH 7.4; diluted with MilliQ-Water from 10x TBS, Bioland Scientific Cat. TBS01-03) three times (5 minutes each) and Tris-A solution (0.1% Triton-X, FisherSci Cat. 85111, in 1x TBS, pH 7.4) at rt for 10 minutes. Antigen retrieval for tissues from synaptophysin staining was facilitated by incubating in citrate buffer at 10 mM with 0.05% Tween-20 (pH 6.0, Sigma-Aldrich Cat. P4922 and Cat. 655204, respectively) at 70$^{o}$C for 30 minutes and recovery in borate buffer (100mM, pH 8.5, Sigma-Aldrich Cat. B0394) at rt for 10 minutes and washed with 1x TBS (2 washes, 5 minutes each). Prior to staining with primary antibodies, sections were incubated in blocking solution with each respective secondary antibody host. This included 3% normal goat serum, NGS with 1% Bovine Serum Albumin or BSA in Tris-A buffer for synaptophysin; 4% BSA in Tris-

A buffer for PSD95; 3% NGS in Tris-A buffer for CD88 (C5aR1) and CD68/IBA1 markers (NGS, Jackson ImmunoResearch Cat. 005-000-121; BSA, Sigma-Aldrich Cat. 05470) at rt for 45 minutes. Sections were incubated in primary antibodies at 4ºC overnight (1:1000 for mouse anti-synaptophysin, Sigma Cat. S5768, in 3% NGS and 1% BSA in Tris-A buffer); 1:1000 for mouse anti-PSD95 (Fisher Scientific Cat. MA1-045, with 2% BSA in Tris-A buffer); 1:500, rat anti-mouse CD88 (FisherSci Cat. MA1-81761), combined with 1:500 rabbit anti-IBA1 (FUJIFILM Wako Cat. 019-19741, with 3% NGS in Tris-A) for CD88-IBA1 dual stain. Sections were placed at room temperature for 1 hour the next day before staining for secondary antibodies. Sections were washed with 1x TBS (3 times, 5 minutes each) and then transferred to secondary antibodies to incubate at room temperature for one hour to visualize the target antigens (1:1000 Goat anti-mouse Alexa Fluor (AF) 647, Abcam Cat. ab150115, for both synaptophysin and PSD95 markers;1:750 goat anti-rat AF 647, Abcam Cat. ab150159, for C5aR1 or CD88 marker, and 1:500 goat anti-rabbit AF 488, Fisher Scientific Cat. a11008, for IBA1 marker). Lastly, brain sections were counterstained with DAPI nuclear dye (1µmol/L, FisherSci Cat. D1306) in 1x TBS at room temperature for 15 minutes. Stained sections were then washed with 1x TBS (2 times, 5 minutes each) and mounted on superfrost slides (FisherSci Cat. 22-037-246) using VectaShield antifade mounting medium (VectaShield Cat. H-1000-10). Staining for CD163 and GFAP followed the similar steps as synaptophysin as described above with a minor change of buffer use (1x PBS, FisherSci) and antibodies. Primary antibodies comprised 1:500 rabbit anti-GFAP (Covance Cat. PRB-571C-100) and 1:200 rabbit anti-CD163 (Abcam, AB182422). The secondary antibodies were 1:500 goat anti-rabbit AF 488 and AF 647, respectively (FisherSci Cat., RRID: AB_143165, RRID: AB_2535864).

Immunohistochemistry staining for CD68-IBA 1 markers follow similar steps above with some minor changes. First, brain sections were washed with PBS with 0.3% Tween 20 (Sigma-Aldrich) three times (5 minutes each) and then incubated in 3% hydrogen peroxide solution (FisherSci Cat. H325-500), with 1% Methanol (FisherSci Cat. A412500) in 1x PBS (FisherSci) at room temperature for 30 minutes to block any endogenous peroxidase activity. Sections were washed in 1x PBS (3 times, 5 minutes each) prior to incubating in blocking solution (4% BSA in PBS with 0.33% Tween-20, Sigma-Aldrich) at room temperature for 30 minutes. Brain sections were then incubated in primary antibodies at 4ºC overnight (1:500, rat anti-mouse CD68, Bio-rad Cat. MCA1957, combined with 1:500 rabbit anti IBA-1, FUJIFILM Wako Cat. 019-1974) with 1% BSA in PBS with 0.3% Tween-20 buffer. Sections were washed and stained in secondary antibodies at room temperature for one hour (1:1000, goat anti-rat AF 647, Abcam Cat. ab150159, for CD68 marker, and 1:500, goat anti-rabbit AF 488, FisherSci Cat. a11008, for IBA1 marker).

Sections were washed and counterstained with DAPI nuclear dye (1µmol/L, FisherSci Cat. D1306) and mounted on superfrost slides (FisherSci Cat. 22-037-246) using VectaShield antifade mounting medium (VectaShield Cat. H-1000-10).

**Confocal microscopy and 3D algorithm-based volumetric quantification**

Immunostained brain sections were imaged at high resolution (1024p) at 0.5 µm thick z stacks using a laser-scanning confocal microscope (Nikon AX) with a 40x oil-immersion objective lens. High-resolution images were deconvoluted, and 3D volume surfaces were created for each antigen of interest using Imaris, an image analysis software (Oxford Instruments). For details, see PMID 33323383. For synaptophysin and PSD95, the surface volumes of synaptic puncta were obtained and analyzed. For CD68/IBA1 and C5aR1/IBA1, co-localization volumetric between the surfaces of two markers for each stain was determined and analyzed. All immunostained image analyses were conducted using automated batch processing, which uniformly applied the parameters to all images to obtain unbiased analyses.

### STATISTICAL ANALYSIS

Statistical analyses were performed to confirm overall significance (GraphPad Prism, v10.0, RRID: SCR_002798). For the analysis of C5aR KO, PMX205 treatment, and cranial irradiation, two-way ANOVA or repeated measures ANOVA, and recommended multiple comparisons tests (Tukey’s, Bonferroni’s) were performed. For the tumor studies, P values were derived from the Mann-Whitney *U* test or long-rank test for survival studies. All results are expressed as the mean values ± SEM. All analyses considered a value of $P \leq 0.05$ to be statistically significant.

| REAGENT or RESOURCES | SOURCE | IDENTIFIER |
| --- | --- | --- |
| **Antibodies** | | |
| FITC anti-mouse CD45 | Biolegend | Cat. 157214; RRID: AB_2894427 |
| PE anti-mouse CD11b | Biolegend | Cat. 101208; RRID: AB_312791 |
| APC anti-mouse CD88 (C5aR1) | Biolegend | Cat. 135808; RRID: AB_10896758 |
| FITC anti-mouse/human CD11b | Biolegend | Cat. 101206; RRID: AB_312788 |
| PE anti-mouse Ly-6C | Biolegend | Cat. 128008; RRID: AB_1186132 |
| Rat anti-Mouse CD68 | Bio-Rad | Cat. MCA1957; RRID: AB_322219 |
| Rabbit anti IBA1 | FUJIFILM Wako | Cat. 019-19741; RRID: AB_839504 |
| Rat α Mouse CD88 | Covance | Cat. MA1-81761; RRID: AB_929568 |
| Rabbit anti-GFAP | Covance | Cat. PRB-571C-100; RRID: AB_291696 |
| Rabbit anti-CD163 | Abcam | Cat. ab182422; RRID: AB_2753196 |
| Mouse anti-Synaptophysin | Sigma-Aldrich | Cat. S5768; RRID: AB_477523 |
| Mouse anti-PSD-95 | FisherSci | Cat. MA1-045; RRID: AB_325399 |
| Goat anti-Rat AF 647 | Abcam | Cat. ab150159; RRID: AB_2566823 |
| Goat anti Rabbit AF 488 | FisherSci | Cat. a11008; RRID: AB_143165 |
| Goat anti Mouse AF 647 | Abcam | Cat. ab150115; RRID: AB_2687948 |
| Donkey anti-Goat AF 647 | FisherSci | Cat. A21447; RRID: AB_2535864 |
| Donkey anti-Rabbit AF 488 | FisherSci | Cat. A21206; RRID: AB_2535792 |
| Donkey anti-Rabbit AF 568 | FisherSci | Cat. A10042; RRID: AB_2534017 |
| Normal Goat Serum | Jackson ImmunoReasearch | Cat. 005-000-121; RRID: AB_2336990 |
| Normal Donkey Serum | Jackson ImmunoReasearch | Cat. 017-000-121; RRID: AB_2337258 |
| DAPI | FisherSci | Cat. D1306; RRID: AB_2629482 |
| **Experimental Model: Cell lines** | | |
| CT2A-Luc astrocytoma cells | Sigma-Aldrich | Cat. SCC195; RRID:CVCL_ZJ60 |
| GL261 glioblastoma cells | NCI-DTP | Cat. Glioma 261; RRID:CVCL_Y003 |
| Contd. | | |

| | | |
|---|---|---|
| **Experimental Model: Organisms/strains** | | |
| C57BL/6J mice | Jackson Labs | Cat. 000664 RRID: IMSR_JAX:000664 |
| C5aR -/- mice | Dr. Rick A. Wetsel, University of Texas Health Science Center, Houston | PMID: 18063050 |
| **Chemicals, peptides, and recombinant proteins** | | |
| Gibco™ DMEM (1X) + GlutaMAX™-I | FisherSci | Cat. 10-569-010 |
| Gibco™ FBS | FisherSci | Cat. 10-082-147 |
| Gibco™ Hibernate™-A Medium | FisherSci | Cat. A1247501 |
| Gibco™ TrypLE™ Express Enzyme | FisherSci | Cat. 12-605-010 |
| Isospire™ (isoflurane) Inhalation Anesthetic | Dechra | N/A |
| Betadine® Surgical Scrub | Patterson Veterinary | Cat. 07-808-8897 |
| Hydrogen Peroxide, 30% (Certified ACS) | FisherSci | Cat. H325-500 |
| Normal Saline (Sodium Chloride) 0.9% | Patterson Veterinary | Cat. 07-800-9424 |
| Buprenorphine HCl (0.3 mg/ml) | MWI Health | N/A |
| VECTASHIELD® Antifade Mounting Medium | VectaShield | Cat. H-1000-10 |
| D-Luciferin, sodium salt | GoldBio | Cat. eLUCNA |
| Gibco™ DPBS (no magnesium, no calcium) | FisherSci | Cat. 14-190-144 |
| Iohexol (Omnipaque™) 350 mgI/ml | GE Healthcare | N/A |
| PMX205 | Trent Woodruff, The University of Queensland | N/A |
| Dimethylsulfoxide (DMSO) | FisherSci | AAJ66650K2 |
| Virkon, premixed spray bottle | University Laboratory Animal Resources | N/A |
| Sodium Chloride, NaCl | Sigma-Aldrich | S9888 |
| Heparin sodium salt from porcine intestinal mucosa | Sigma-Aldrich | Cat. H3149 |
| Gibco™ HBSS (no magnesium, no calcium) | FisherSci | Cat. 14-175-079 |
| Collagenase from Clostridium histolyticum | Sigma-Aldrich | Cat. C0130-500MG |
| Nα-Tosyl-L-lysine chloromethyl ketone hydrochloride | Sigma-Aldrich | Cat. T7254-100MG |
| DNAse 1 | Zymo | Cat. E1011-A |
| Gibco™ HEPES (1M) | FisherSci | Cat. 15-630-080 |
| Gibco™ RPMI 1640 Medium, GlutaMAX™ Supplement | FisherSci | Cat. 61870036 |
| Cytiva Percoll™ Centrifugation Media | FisherSci | Cat. 45-001-748 |
| Gibco™ DPBS (10X), no calcium, no magnesium | FisherSci | Cat. 14-200-075 |
| GenClone 25-508 DPBS, 1X, without Ca, Mg, Phenol Red, 0.1um Sterile Filtered | Genesee Scientific | Cat. 25–508 |
| Invitrogen™ eBioscience™ Flow Cytometry Staining Buffer | FisherSci | Cat. 501128916 |
| Paraformaldehyde | Sigma-Aldrich | Cat. 158127 |
| Sodium azide | Sigma-Aldrich | Cat. S2002 |
| Sucrose | Sigma-Aldrich | Cat. S7903 |
| Tissue-Tek* O.C.T. Compound | VWR | Cat. 25608-930 |
| Tris-buffered Salin (10X TBS) | Bioland Scientific | Cat. TBS01-03 |
| Triton™ X-100 Surfact-Amps™ Detergent Solution | FisherSci | Cat. 85111 |
| Phosphate-Citrate Buffer with Sodium Perborate | Sigma-Aldrich | Cat. P4922 |

| | | |
|---|---|---|
| TWEEN® 20 | Sigma-Aldrich | Cat. 655204 |
| Boric acid | Sigma-Aldrich | Cat. B0394 |
| Bovine Serum Albumin | Sigma-Aldrich | Cat. 05470 |
| QIAzol Lysis Reagent (200ml) | Qiagen | Cat. 79306 |
| M-MLV Reverse Transcriptase | Promega | Cat. M170A |
| RT 5X buffer | Promega | Cat. M531A |
| Oligo (dT) 18 Primer | FisherSci | Cat. S0131 |
| dNTP Set (100 mM) | FisherSci | Cat. 10297018 |
| RNasin® Ribonuclease Inhibitor | Promega | Cat. N2511 |
| DL-Dithiothreitol (DTT) | FisherSci | Cat. 707265ML |
| DEPC-Treated Water | FisherSci | Cat. AM9920 |
| **Critical commercial assays** | | |
| RNeasy Plus Universal Mini Kit | Qiagen | Cat. 73404 |
| Applied Biosystems™ TaqMan™ Universal PCR Master Mix, no AmpErase™ UNG | FisherSci | Cat. 4324018 |
| TaqMan Gene Expression Assays (Assays ID: GAPDH Mm99999915_g1) | FisherSci | Cat. 4331182 |
| TaqMan Gene Expression Assays (Assays ID: CD88 Mm00500292_s1) | FisherSci | Cat. 4331182 |
| Nanostring nCounter Mouse Neuroinflammation | Nanostring | Cat. XT-CSO-MNROI1-12 |
| **Software and algorithms** | | |
| Living Image Software | Revvity | Cat.128110 |
| FlowJo | BD Biosciences | N/A |
| OsiriX Lite Software | Pixmeo SARL | N/A |
| Noldus EthoVisionXT 17 Tracking Software | Noldus | N/A |
| FreezeFrame | Coulbourn Instruments | N/A |
| Imaris 10.0 | Oxford Instruments | N/A |
| nSolver software version 4.0 | Nanostring | N/A |
| nSolver Advanced Analysis | Nanostring | N/A |
| Advanced Treatment Planning | Precision X-Ray | N/A |
| **Other** | | |
| RC2 – Rodent Circuit Controller | VetEquip | Cat. 922100 |
| Ultra Precise Digital Just for Mouse Stereotaxic Instrument | Stoelting | Cat. 51730UD |
| Sterile Alcohol Prep Pads | FisherSci | Cat. 06-669-62 |
| Feather™ Single-Use Scalpels | FisherSci | Cat. 08-927-5B |
| 10 µL Microliter Syringe Model 701 N | Hamilton | Cat. 80300 |
| Pivetal® WebCryl™ Sutures | Patterson Veterinary | Cat. 07-891-0161 |
| 3M™ Vetbond™ Tissue Adhesive Bottle | Patterson Veterinary | Cat. 07-805-5031 |
| Fisherbrand™ Superfrost™ Plus Microscope Slides | FisherSci | Cat. 22-037-246 |
| IVIS Lumina III *in vivo* Imaging System | Revvity | Cat. CLS136334 |
| SmART+ irradiator | Precision X-Ray | N/A |
| Noldus EthoVisionXT 17 Video Capture Hardware System | Noldus | N/A |
| C1000 Touch™ Thermal Cycler with Dual 48/48 Fast Reaction Module | Bio-Rad | Cat. 1851148 |
| Nanodrop | FisherSci | Cat. 13-400-525 |
| Bioanalyzer 2100 | Agilent Technologies | N/A |
